# Historical contingency shapes adaptive radiation in Antarctic fishes

**DOI:** 10.1101/478842

**Authors:** Jacob M. Daane, Alex Dornburg, Patrick Smits, Daniel J. MacGuigan, M. Brent Hawkins, Thomas J. Near, H. William Detrich, Matthew P. Harris

**Affiliations:** Department of Marine and Environmental Sciences, Northeastern University Marine Science Center, Nahant, MA 01908, USA; North Carolina Museum of Natural Sciences, Raleigh, NC 27601, USA; Orthopaedic Research Laboratories, Department of Orthopaedic Surgery, Boston Children’s Hospital, Boston, MA 02115, USA; Department of Ecology and Evolutionary Biology, Yale University, New Haven, CT 06520, USA; Museum of Comparative Zoology, Harvard University, Cambridge, MA 02138, USA; Department of Genetics, Harvard Medical School, Boston, MA 02115, USA; Peabody Museum of Natural History, Yale University, New Haven, CT 06520, USA

## Abstract

Adaptive radiation illustrates the links between ecological opportunity, natural selection, and the generation of biodiversity (1). Central to adaptive radiation is the association between a diversifying lineage and the evolution of key traits that facilitate the utilization of novel environments or resources (2, 3). However, is not clear whether adaptive evolution or historical contingency is more important for the origin of key phenotypic traits in adaptive radiation (4, 5). Here we use targeted sequencing of >250,000 loci across 46 species to examine hypotheses concerning the origin and diversification of key traits in the adaptive radiation of Antarctic notothenioid fishes. Contrary to expectations of adaptive evolution, we show that notothenioids experienced a punctuated burst of genomic diversification and evolved key skeletal modifications before the onset of polar conditions in the Southern Ocean. We show that diversifying selection in pathways associated with human skeletal dysplasias facilitates ecologically important variation in buoyancy among Antarctic notothenioid species, and demonstrate the sufficiency of altered *trip11, col1a2* and *col1a1* function in zebrafish (*Danio rerio*) to phenocopy skeletal reduction in Antarctic notothenioids. Rather than adaptation being driven by the cooling of the Antarctic (6), our results highlight the role of exaptation and historical contingency in shaping the adaptive radiation of notothenioids. Understanding the historical and environmental context for the origin of key traits in adaptive radiations provides context in forecasting the effects of climate change on the stability and evolvability of natural populations.

## Main Text

During adaptive radiation, ecological opportunity and key phenotypic traits interact to facilitate the expansion of populations into new niches (1). Deciphering the genotype-phenotype relationships of these key traits provides important insight into the historical circumstances that result in phenotypic diversification (7, 8). While investigations into the genetic basis of key adaptive traits have been limited by focusing primarily on single species (9), advances in sequencing capabilities now allow for the efficient collection of genomic sequence data for scores of species. This potential for increased taxonomic, genome-wide sampling provides opportunities to investigate the macroevolutionary mechanisms driving evolution across clades. For example, a major question in the study of adative radiation is the relative role for adaptation and exaptation in facilitating the diversification of species. Adaptive evolution can result from ecological opportunities, such as those brought about by environmental changes, driving the origin of key innovations and accelerating lineage diversification (2, 3). In contrast, exaptation suggests that trait novelty may arise at any time, and that adaptive change is merely the consequence of a shift in selection that enables lineages to opportunistically explore new ecospace using a trait that had previously evolved under different circumstances (10). These two hypotheses have long been central in the debate concerning the role of punctuated versus gradualistic evolution in the generation of biodiversity (10, 11). Here, we establish the genetic basis of ecologically important traits, as well as their phylogenetic origin in a species-rich adaptive radiation. Through consideration of closely related lineages that are not a part of the adaptive radiation, we can specifically investigate the relative importance of adaptive evolution and contingency in adaptive radiation.

To test hypotheses concerning the origin of traits that facilitate adaptive radiation, we focused on Antarctic notothenioids (Cryonotothenioidea), an iconic example of adaptive radiation in marine vertebrates (6, 12). The diversification of cryonotothenioids followed the progressive cooling of Antarctica that initiated ~33 million years ago (6), and was coincident with the extinction of a phylogenetically diverse and cosmopolitan fish fauna (13). This combination of climate change and vacated niches presented notothenioids with the opportunity to diversify into a large range of benthic and water column habitats. All notothenioids lack a swim bladder, the primary buoyancy organ in most teleost fishes; however, there are substantial differences in buoyancy among notothenioid species that are correlated with habitat and resource utilization (6, 13). These differences in buoyancy are achieved through the reduction of skeletal density coupled with the accumulation of corporeal lipids that provide static lift (13–15). In the narrative of the Antarctic cryonotothenioid adaptive radiation, these ecologically important traits are hypothesized to have arisen during the onset of polar conditions; however, the evolutionary origin of reduced skeletal density in the context of early-diverging non-Antarctic notothenioid lineages challenges this scenario (15).

To develop a comprehensive phylogenomic perspective on the Antarctic notothenioid adaptive radiation, we combined sequence information from the genome of *Notothenia coriiceps* and select other percomorph teleost species to design a comparative DNA probe set for systematic targeted enrichment of ~250,000 coding and conserved non-coding (CNE) elements comprising over 40 Mb of genomic sequence (**Fig. S1**). Using this probe set, we captured coding and non-coding sequences from a phylogenetically-rich sampling of 46 notothenioid species that includes all three early-diverging non-Antarctic lineages and two species from the closely related Percidae (**Fig. S1**). We achieved 89-95% coverage of targeted regions for each sampled species (**Fig. S1; Table S1,S2**), permitting identification of an average of 95,000 fixed species-specific SNPs and 60,000 heterozygous SNPs for each lineage (**Table S3**). As proof of the sensitivity of the dataset, we confirmed known cases of adult hemoglobin gene loss within the icefishes (**Fig. S2**) (16). However, the power of this dataset was further illustrated by identification of previously uncharacterized deletion of two putative embryonic hemoglobin genes (**Fig. S2**). Thus, this approach provides a robust and efficient method by which to characterize variation across taxonomically rich clades, even when separated by significant evolutionary distances.

Given expectations from the adaptive radiation of East African cichlids (17), we tested whether shifts in the rate of nucleotide evolution correspond to the onset of rapid lineage diversification in the cryonotothenioid radiation. Using a time-calibrated phylogeny of most living notothenioids and a Bayesian framework to assess shifts in speciation rates (12, 18), we confirmed a shift in lineage diversification at the origin of the antifreeze-bearing Antarctic cryonotothenioids and an additional acceleration in diversification within the Plunderfishes (Artedidraconidae) (Fig. 1A) (6). Intriguingly, these shifts do not correspond with accelerated rates of nucleotide evolution. Instead, we found that the majority of extant cryonotothenioid sequence diversity is derived from a period of significantly high rates of genomic evolution that occurred in the ancestral lineage that includes the most recent common ancestor of the non-Antarctic distributed *Eleginops maclovinus* and the Antarctic cryonotothenioids (Eleginopsioidea, Fig. 1B, **Table S4**). This demonstrates that a major change in molecular rate variation preceded the onset of global cooling and adaptive radiation of cryonotothenioids by well over 10 million years (Fig. 1C). An assessment of the accumulation of genomic divergence through relaxed molecular clock models (Fig 1D), along with an analysis of synonymous (dS) and nonsynonymous substitution (dN) rates (Fig. 1E), further substantiates a high rate of nucleotide evolution prior to the origin and diversification of cryonotothenioids. This acceleration of overall genomic change may have been critical for accumulating the genetic diversity that provided the substrate for phenotypic diversification during the early radiation of cryonotothenioids.

**Fig. 1.**
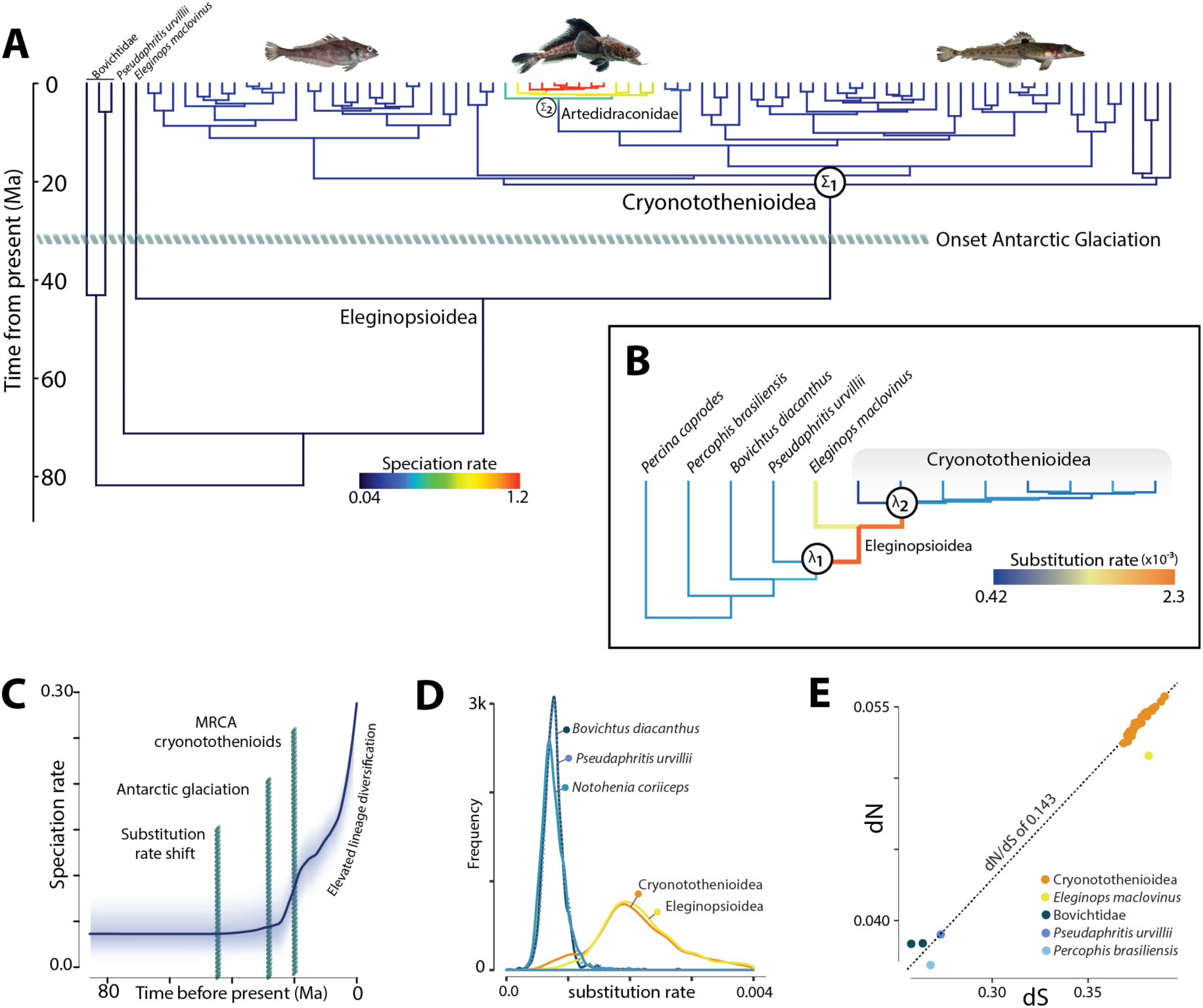
Punctuated elevation in genomic diversification prior to ecological change and adaptive radiation. (**A**) Bayesian analysis of lineage diversification rate on time-calibrated notothenioid phylogeny (12, 18). ∑ indicates change in speciation rate. Colors on branches correspond to the mean of the marginal posterior density of estimated speciation rates, with a shift from low (0.04 lineages/Ma) to faster rates of speciation (0.046 lineages/Ma) at the onset of the notothenioid adaptive radiation. (**B**) Relaxed molecular clock model of nucleotide substiution rate (substitutions/site/Ma) during notothenioid evolution revealing a transient elevation in substitution rate prior to the increase in species diversification. Substitution rates estimated from 1,062 independent gene trees constructed using the random local clock model in BEAST2; λ indicates change in substitution rate. Colors on branches correspond to mean substitution rate, from low (0.00042 substitutions/site/Ma) to the high (0.00228 substiutions/site/Ma) observed on the branch leading to *Eleginops maclovinus* and all cryonotothenioids. (**C**) Elevation in speciation rate well after increase in nucleotide substitution rate and climate change events that precipitated adaptive diversification. Shading indicates 10-90% Bayesian credible region across time. (**D**) Distribution of substitution rates from select species in (**B**). (**E**) Average synonymous (dS) and non-synonymous (dN) substitution rate from over 4,000 pairwise alignments for each species relative to the outgroup, *Percina caprodes*.

As buoyancy adaptations are key traits that facilitated notothenioid diversification into the water column during the adaptive radiation, we assessed patterns of skeletal density throughout the phylogeny using computerized tomography (Fig. 2A). We find that, in parallel with the observed increase in nucleotide substitution rate, a broad reduction in bone density occurred prior to the recent common ancestor of *Eleginops maclovinus* (19) and the Antarctic cryonotothenioids (Eleginopsioidea, Fig. 2A) and was retained throughout the clade. Consistent with these decreases in bone density, we found biased diversifying selection in genes associated with human skeletal dysplasias and mineralization defects in Eleginopsioidea relative to the well-ossified sister lineage, *Pseudaphritis urvillii* (Fig. 2B,C). This evolutionary reconstruction is contrary to the expectation of key innovation-driven diversification, but rather indicates that prior to the onset of polar conditions notothenioid lineages were beginning to utilize habitats in the water column through a modification of buoyancy that was enabled through a derived reduction in bone density.

**Fig. 2.**
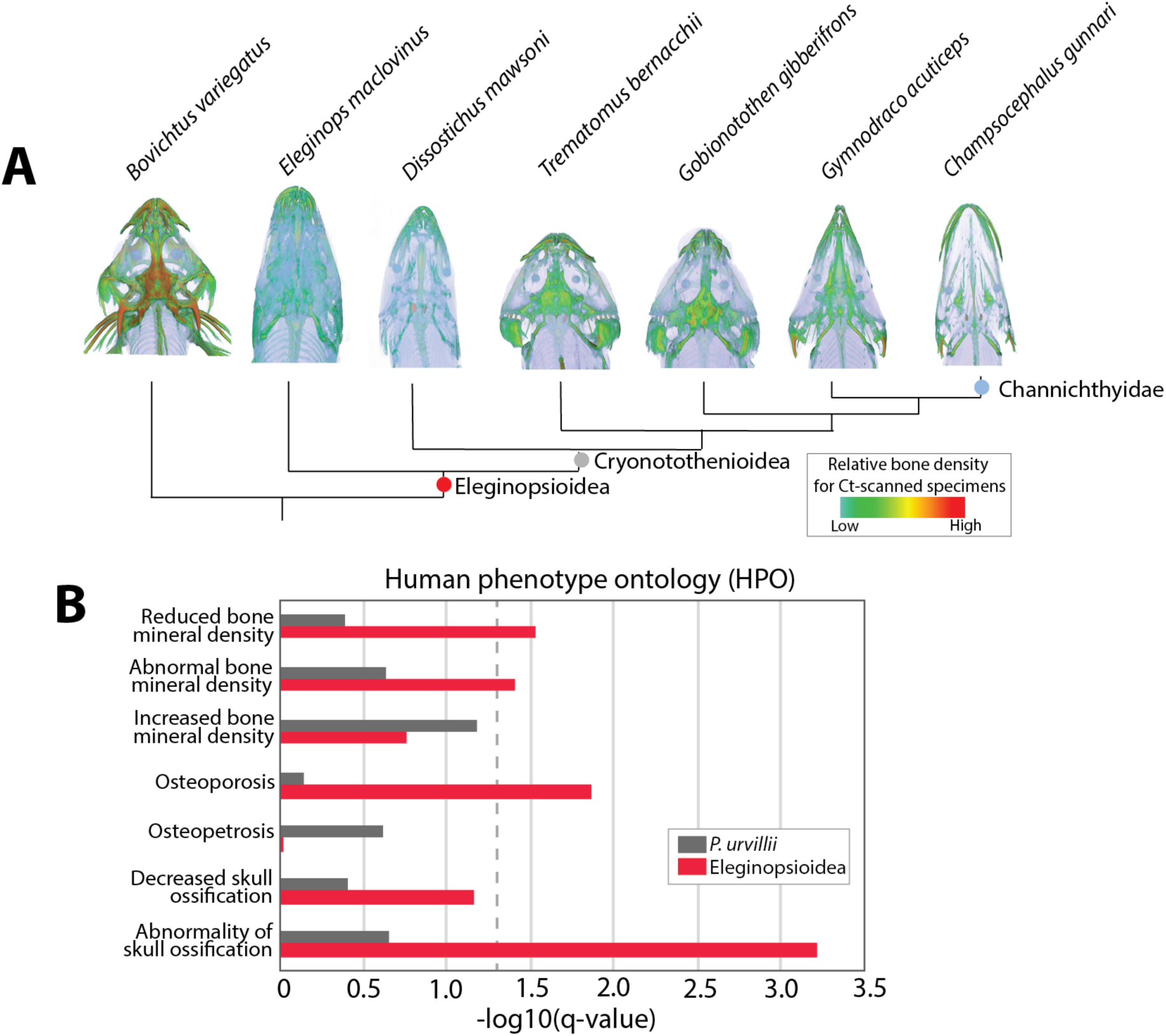
Skeletal reduction as an exaptation for buoyancy in the cryonotothenioid radiation. (**A**) Computed tomography (CT) of notothenioid skulls showing relative skeletal density across the phylogeny and decrease in skeletal density prior to cryonotothenioid radiation. (**B**) Comparison of significance of enrichment for polygenic selection on represenative bone-density associated HPO terms between Eleginopsioidea and well-ossified sister group *Pseudaphritis urvillii*. Dashed line indicates FDR q-value of 0.05.

To further explore the genetic basis of the evolved reduction in bone density, we assessed patterns of diversifying selection within the phylogeny. Among *Eleginops maclovinus* and all cryonotothenioids, we identified shared selective signatures in clinically relevant skeletal genes, such as *collagen1a1* and *collagen1a2* (Fig. 3A, **Table S5**). The function of these genes are conserved among disparate lineages of vertebrates (20) and nonsynonymous mutations in these collagens can lead to severe osteogenesis imperfecta in humans (21). Notably, previous studies show that *collagen1* expression is reduced in the developing skeleton of notothenioid embryos, providing further evidence that broad changes at collagen loci are associated with skeletal variation (22). In addition to selection on collagens, we identified diversifying selection in an unlikely gene candidate for skeletal variation, *trip11*(*gmap-210*) (Fig 3A, **Table S5**). This gene is conserved across eukaryotes and functions in vesicle tethering in the cis-Golgi membrane (23). In humans, mutations in *TRIP11* lead to severe skeletal deficiencies that lead to peri-natal lethality (24). Intriguingly, both *collagen1a1a* and *trip11* are also under diversifying selection and/or accelerated sequence evolution within the further diversification of the Channichthyidae, which have evolved an additional and significant reduction in bone density (Fig. 2A; Fig 3A; **Tables S5,S6,S7,S8**) (15). This parallelism suggests constraint in the types of mutations in evolution that can drive changes in skeletal density while maintaining viability.

**Fig. 3.**
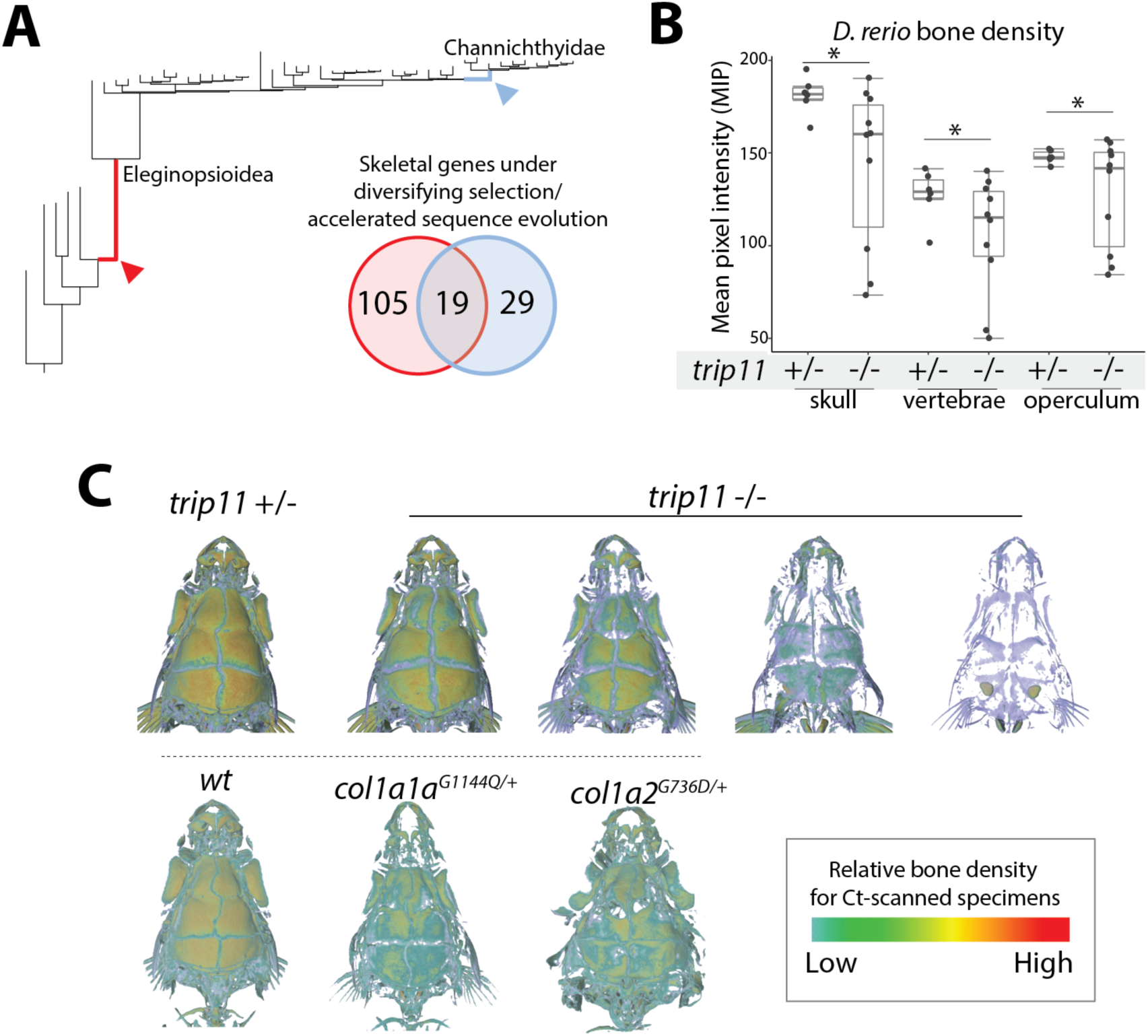
Skeletal genes under diversifying selection uncovers genetic mechanisms regulating bone density. (**A**) Comparison of genes under positive selection (aBSREL, p<0.05) and/or accelerated sequence evolution (phyloP, p<0.05) on the branch leading to *Eleginops maclovinus* and Antarctic cryonotothenioids (Eleginopsioidea) with the branch leading to the icefishes, Channichthyidae. (**B**) Quantitation of skeletal density from µCT analysis of zebrafish mutants in *trip11*. Density is the average pixel intensity. n=6 scanned *trip11^+/-^* and n=10 *trip11^-/-^* individuals. Center line is mean density, box bounds indicate lower and upper quartiles, whiskers extend to a maximum of 1.5 times the interquartile range. (**C**) Micro-computed tomography (µCT) of zebrafish showing reduction in bone density in the cranium of zebrafish mutant models of genes under selection in Eleginopsioidea. Representative µCT images of zebrafish skulls from heterozygous and homozygous *trip11* mutants, showing reduced bone density and variable penetrance. Reduction in density in the cranium caused by mutations in *col1a1a^dmh14/+^* (G1144Q) and *col1a2^dmh15/+^* (G736D). *: one tailed t-test p-value <0.05.

To experimentally test the potential for alterations in *trip11, col1a1* and *col1a2* function to impart non-lethal skeletal phenotypes that are consistent with those observed in notothenioids, we analyzed mutants stemming from genetic screens and CRISPR-Cas9 gene editing in zebrafish (*Danio rerio*). Contrary to expectations from humans, zebrafish homozygous for loss-of-function alleles of *trip11* were viable with no obvious external morphological abnormalities (Fig. 3B,C). However, whereas heterozygous siblings had density patterns comparable to wild-type zebrafish, similarly size and age-matched homozygous *trip11* mutants had significantly reduced skeletal density (Fig. 3B,C). In affected individuals, all bones investigated, including the calvaria, vertebrae and operculum were reduced in density (Fig. 3B,C) and generally phenocopied the pattern observed in adult *Eleginops maclovinus* and cryonotothenioids (Fig. 2A). Notably, the expressivity of the *trip11* skeletal phenotype was variable, which is consistent with the presence of background genetic modifiers. Similarly, we show mutant models of *col1a1^dmh14/+^* (G1144Q) and *col1a2 ^dmh54/+^* (G736D) in zebrafish also cause a reduction in bone density (Fig. 3C) (20). These experiments confirm that changes to the loci under selection yield phenotypes consistent with those observed in *E. maclovinus* and species of the cryonotothenioid radiation.

Our results reveal that historical contingency was a major factor in shaping the adaptive radiation of notothenioids. As the onset of polar conditions ~33 mya decimated the teleost fauna of the Southern Ocean, notothenioids were the only surviving lineage poised to occupy newly open niche space in a range of benthic and water column habitats (25). Rather than being driven by directional selection to evolve extreme phenotypes in response to the onset of polar conditions (26, 27), the genomic substrate for reduced ossifications and buoyancy modifications had long been established and was coopted to facilitate the ecological diversification that characterizes the notothenioid adaptive radiation. These results provide an alternative view on the impact of climate change in driving extreme adaptations in the Southern Ocean (28, 29). As we march further into the Anthropocene, our results caution that as both ecosystems and climate continue to change worldwide, the expectation that rapid adaptation will be dominant factor in predicting the response of biodiversity requires careful consideration (30).

## Supporting information

## Acknowledgements

The authors thank Dr. Erin Snay and Leah Oberg in Department of Nuclear Medicine and Molecular Imaging at Boston Children’s Hosptial for assistance in computed tomography of adult specimens.

## Author contributions

Conceived and designed the study: JMD, HWD, MPH. Performed the experiments: JMD, PS, MBH. Analyzed the data: JMD, AD, TJN, DJM, HWD, MPH. JMD, AD, and TJN wrote the first drafts of the manuscript. All authors contributed to the writing of the final manuscript.

## Data and Materials Availability

Upon acceptance, the assembled contigs and annotations for all species in the dataset will be made available on the Zenodo repository (https://zenodo.org/). Correspondence and requests for materials should be addressed to.

## References

1 D. Schluter, The Ecology of Adaptive Radiation (Oxford University Press, 2000).

2 D. L. Rabosky, Phylogenetic tests for evolutionary innovation: The problematic link between key innovations and exceptional diversification. Philos. Trans. R. Soc. B Biol. Sci. 372 (2017).

3 J. T. Stroud, J. B. Losos, Ecological Opportunity and Adaptive Radiation. Annu. Rev. Ecol. Evol. Syst. 47, 507–532 (2016).

4 J. B. Losos, Lizards in an evolutionary tree: ecology and adaptive radiation of anoles (University of California Press, 2009).

5 S. J. Gould, Wonderful Life: the Burgess Shale and the Nature of History (W. W. Norton, 1989), vol. 17.

6 T. J. Near et al., Ancient climate change, antifreeze, and the evolutionary diversification of Antarctic fishes. Proc. Natl. Acad. Sci. 109, 3434–3439 (2012).

7 Y. F. Chan et al., Adaptive Evolution of Pelvic Reduction of a Pitx1 Enhancer. Science. 327, 302–306 (2010).

8 M. E. Santos et al., The evolution of cichlid fish egg-spots is linked with a cis-regulatory change. Nat. Commun. 5 (2014).

9 F. C. Jones et al., The genomic basis of adaptive evolution in threespine sticklebacks. Nature. 484, 55–61 (2012).

10 S. J. Gould, The Structure of Evolutionary Theory (Harvard University Press, 2002).

11 D. Jablonski, Approaches to Macroevolution: 1. General Concepts and Origin of Variation. Evol. Biol. 44, 427–450 (2017).

12 A. Dornburg, S. Federman, A. D. Lamb, C. D. Jones, T. J. Near, Cradles and museums of Antarctic teleost biodiversity. Nat. Ecol. Evol. (2017).

13 J. T. Eastman, Antarctic Fish Biology: Evolution in a Unique Environment (Academic Press, Inc, San Diego, CA, 1993).

14 A. L. DeVries, J. T. Eastman, Lipid sacs as a buoyancy adaptation in an Antarctic fish. Nature. 271 (1978), pp. 352–353.

15 J. T. Eastman, L. M. Witmer, R. C. Ridgely, K. L. Kuhn, Divergence in skeletal mass and bone morphology in antarctic notothenioid fishes. J. Morphol. 275, 841–61 (2014).

16 T. J. Near, S. K. Parker, H. W. Detrich, A genomic fossil reveals key steps in hemoglobin loss by the Antarctic icefishes. Mol. Biol. Evol. 23, 2008–2016 (2006).

17 D. Brawand et al., The genomic substrate for adaptive radiation in African cichlid fish. Nature (2014).

18 D. L. Rabosky, Automatic detection of key innovations, rate shifts, and diversity-dependence on phylogenetic trees. PLoS One. 9 (2014).

19 J. T. Eastman, L. M. Witmer, R. C. Ridgely, K. L. Kuhn, Divergence in skeletal mass and bone morphology in antarctic notothenioid fishes. J. Morphol. 275, 841–861 (2014).

20 C. Gistelinck et al., Zebrafish type I collagen mutants faithfully recapitulate human type I collagenopathies Short title: Zebrafish mutants mimic type I collagenopathies. PNAS, 1–10 (2018).

21 F. S. Van Dijk, D. O. Sillence, Osteogenesis imperfecta: Clinical diagnosis, nomenclature and severity assessment. Am. J. Med. Genet. Part A. 164, 1470–1481 (2014).

22 R. C. Albertson et al., Molecular pedomorphism underlies craniofacial skeletal evolution in Antarctic notothenioid fishes. BMC Evol. Biol. 10, 4 (2010).

23 T. M. Witkos, M. Lowe, The Golgin Family of Coiled-Coil Tethering Proteins. Front. Cell Dev. Biol. 3, 1–9 (2016).

24 P. Smits et al., Lethal skeletal dysplasia in mice and humans lacking the golgin GMAP-210. N. Engl. J. Med. 362, 206–216 (2010).

25 J. T. Eastman, A. R. McCune, Fishes on the Antarctic continental shelf: evolution of a marine species flock? J. Fish Biol. 57, 84–102 (2000).

26 L. Chen, A. DeVries, C. Cheng, Evolution of antifreeze glycoprotein gene from a trypsinogen gene in Antarctic notothenioid fish. Proc. Natl. Acad. Sci. 94, 3811–3816 (1997).

27 Z. Chen et al., Transcriptomic and genomic evolution under constant cold in Antarctic notothenioid fish. Proc. Natl. Acad. Sci. U. S. A. 105, 12944–12949 (2008).

28 S. L. Chown et al., The changing form of Antarctic biodiversity. Nature. 522, 431–438 (2015).

29 S. L. Chown et al., Antarctica and the strategic plan for biodiversity. PLoS Biol. 15, 1–10 (2017).

30 K. T. Bilyk, L. Vargas-Chacoff, C. H. C. Cheng, Evolution in chronic cold: Varied loss of cellular response to heat in Antarctic notothenioid fish. BMC Evol. Biol. 18, 1–16 (2018).

