## Supplementary material for "Historical contingency shapes adaptive radiation in Antarctic fishes"

**This PDF file includes:**

Materials and Methods

Figs. S1 to S8

Tables S1 to S11

Supplemental References

### Table of Contents


**Materials and Methods** 4

**1. Data generation** 4
1.1. Custom targeted sequence enrichment design 4
1.2. Sample preparation and sequencing 4
1.3. Targeted sequence enrichment and next generation sequencing 5
**2. Reference contig assembly** 5
2.1. Processing of sequencing reads 5
2.2. read binning into orthology groups by BLAST 5
2.3. *de novo* contig assembly 6
2.4. Contig merging and filtering 6
**3. Analysis of assembled reference contigs** 7
3.1 Comparison of assembled contigs to reference genome 7
3.2. Estimation of read coverage and depth of targeted regions 7
3.3. Distribution of read coverage across the dataset 7
3.4. Recovery of population variation 7
**4. Identification of orthologs** 8
4.1. Treatment of exons with a predicted history of duplication 8
4.2. Identification of orthologous sequences 8
4.2. Simulation of ortholog identification approach 8
**5. Processing of reference contigs for comparative genomics** 8
5.1. Multiple sequence alignment 8
5.2. Reconstruction of gene sequences from exon data 9
**6. Comparative evolutionary analysis of genetic data** 9
6.1. Notothenioid phylogeny 9
6.2. Quantification of lineage diversification dynamics 9
6.3. dS and dN estimates of substitution rate 10
6.4. Molecular clock models of substitution rate 10
6.5. Detection of diversifying selection 10
6.6. Ontology enrichment 10
**7. Experimental analysis of bone density** 11
7.1. Zebrafish husbandry and genetic lines 11
7.2. Zebrafish genome editing 11
7.3. Analysis of skeletal density through computed tomography (CT) 11

**Figures** 13
Fig. S1 13
Fig. S2 14
Fig. S3 15
Fig. S4 16
Fig. S5 17
Fig. S6 18
Fig. S7 19
Fig. S8 20

**Tables** 21
Table S1 21
Table S2 23
Table S3 25
Table S4 27
Table S5 28
Table S6 31
Table S7 33
Table S8 34
Table S9 35
Table S10 41
Table S11 43

**Supplemental References** 44

### Materials and Methods

#### 1. Data generation

##### 1.1. Custom targeted sequence enrichment design

We based the design of the DNA enrichment baits primarily on the *Notothenia coriiceps* genome, the most closely related species to those targeted for sequencing with a published reference assembly (*1*). To account for regions that are unannotated, under drift, or not easily identified within the *N. coriiceps* assembly, we included targeted regions from several outgroup genomes as detailed below. By having the same element potentially represented by more than one genome, this strategy allowed us to mitigate against genome assembly and annotation artifacts while facilitating hybridization of diverse species to the capture probes. Elements were identified in the *N. coriiceps* genome using BLASTN (ncbi-blast-2.2.30+ ; -max_target_seqs 1 -outfmt 6). If the BLASTN hit had a E-value < 0.0001 and covered >80% of the query sequence, we included this region from the *N. coriiceps* genome. If the region was not identified, or had <85% identity in *N. coriiceps*, we retained the version from the genome of origin.

Coding exons were identified from annotations of the *Notothenia coriiceps* (*1*), stickleback (*Gasterosteus aculeatus*; BROADS1), and European sea bass (*Dicentrachus labrax*) genomes (*2*). Conserved non-coding elements (CNEs) were defined from the constrained elements identified in the stickleback and tilapia (*Oreochromis niloticus*; Orenil1.0) genomes from the Ensembl compara 11-way teleost whole genome alignment (*3*). We also included predicted miRNA hairpins from miRbase (*4*) and ultraconservative non-coding (UCNE) elements from UCNEbase (*5*). miRNA hairpins were padded to be >100bp. CNEs, miRNAs, and UCNEs that overlapped coding exons were removed using Bedtools (v2.26.0) intersectBed (*6*), and CNEs <100bp were additionally excluded to facilitate space in the sequence capture design. Where the constrained regions that defined the CNEs overlapped with annotations pertaining to specific miRNAs and UCNEs, the latter annotations were prioritized.

Targeted elements were submitted to Nimblegen for final probe design and the manufacturing of a Nimblegen SeqCap EZ Developer Library (Roche cat 06471684001) had 63,838,670 bp of capture space targeting 318,929 elements from four reference genomes (88.9% *Notothenia coriiceps*, 7.7% *Gasterosteus aculeatus*, 3.0% *Dicentrarchus labrax*, 0.4% *Oreochromis nicoleatus*). Accounting for redundancy of orthologous target regions between the genomes, the final design targeted 258,176 unique elements, of which 206,503 were predicted protein coding exons and 51,673 constrained non-coding regions with 85.0% coverage of targets not found in the *N. coriiceps* reference genome (47,097 elements; **Table S2**).

##### 1.2. Sample preparation and sequencing

Frozen tissue samples were acquired from the HWD and TJN labs, and from the Yale Peabody Museum frozen tissue collection (**Table S9**). DNA from each species was isolated using Qiagen DNeasy Blood and Tissue kits, sequencing multiple individuals per species to account for population variation. For each species, equal amounts of DNA from each individual were pooled prior to shearing and library preparation. The pooled-population DNA was then sheared to an average size of 200bp using the Covaris E220 ultrasonicator in 130ul Covaris microTUBEs (Duty

Cycle: 10%, Intensity: 5, Cycles/Burst: 200, Time: 300s, Temp: 4°C). Shearing was performed in shearing buffer: 10mM Tris, 0.1mM EDTA, pH8.3.

1.3. Targeted sequence enrichment and next generation sequencing

Sequencing libraries were constructed from 1µg of DNA using KAPA Library Prep kit (Roche cat# 07137923001), following standard protocol with barcoding and dual-SPRI size selection to generate libraries between 200-450bp. Sequencing libraries were hybridized, recovered, and amplified according to the standard protocol (Nimblegen SeqCap EZ Library SR User's Guide v4.3) with the following changes: hybridization was performed at 45°C instead of 47°C to allow for more mismatches between sequencing libraries and probes, and we used SeqCap Developer Reagent (Roche cat# 06684335001) instead of Human CotI DNA to block non-specific hybridization as recommended by the protocol. Since the majority of the capture probes were designed based on the sequence of a *Nothothenia coriiceps*, we hybridized species in groups to limit potential competition between sequencing libraries of varied relatedness to the capture baits. Captured libraries were pooled for 100bp single-end sequencing using Illumina HiSeq 2500. We targeted multiplexing of 8-9 species per HiSeq 2500 flow cell, totaling six flow cells.

**2. Reference contig assembly**

The contig assembly approach is modified from a previously defined pipeline for cross-species targeted sequence enrichment datasets (*7*). Briefly, sequencing reads are grouped into bins by homology to a targeted element (exon, CNE, etc.) and then assembled into contigs *de novo* (**Fig. S3)**

2.1. Processing of sequencing reads

Prior to contig assembly, low quality bases within sequencing reads were masked using the FASTX-Toolkit (fastq_masker; -Q 33) (http://hannonlab.cshl.edu/fastx_toolkit). Illumina adapter sequences were then trimmed using Trimmomatic v0.36 (*8*). Identical sequencing reads were then collapsed using the fastx toolkit v0.0.13 (fastx_collapser; -Q 33).

2.2. Read binning into orthology groups by BLAST

Reads were grouped by homology before contig assembly, using both blastn and dc-megablast (v2.6.0+, -max_target_seqs 2 -outfmt 6). This dual-BLAST approach accounts for variation between sequencing reads and the reference genome from which the sequencing baits were defined (*9*). As short target exons and CNEs can produce disproportionately small blastn E-values, we used an adaptive E-value cutoff based on the size of the target region. For target regions >25 bp, the cutoff was E-value ≤1e-05. For targets that were ≤25 bp the E-value cutoff was ≤1e-04 and for targets that were ≤20bp the E-value cutoff was ≤1e-03. Reads were further excluded if a substantial portion of the read (>10bp) overlapped the target interval without being included in a blastn hit. The best resulting E-value from either blastn or dc-megablast was selected, with blastn selected in the event of a tie. As dc-megablast utilizes a mismatch-tolerant seed template, inclusion of dc-megablast resulted in the additional recovery of 30,000-50,000 sequencing reads per species (out of an average of 20,000,000 with total blastn hits) and the assembly of 50-150 more target regions than would be assembled by blastn alone.

2.3. *de novo* contig assembly

CAP3 was used to assemble contigs *de novo* from the bins of reads that have high homology to specific target regions (i.e. the same exon, CNE, etc.) that were identified by BLAST (*10*). Reads were reverse complemented if necessary in order to put everything into the same complement strand as the target region from the reference genome. To accelerate CAP3 assembly, overlapping reads were first merged into smaller contigs using Usearch and then mixed with original reads as input for CAP3 (-id 0.97 -fastq_maxdiffs 3 -fastq_minovlen 5). For CAP3 assembly, we required a minimum read overlap of 16bp and 96% identity between reads during contig assembly (-o 16 -p 96). This cutoff has an effect of separating the reads stemming from duplication events into separate contigs as long as there is >4-6% variance between the paralogous regions.

We simulated the ability of this pipeline to distinguish copy number variants (**Fig. S4A**), generating a 300bp random DNA sequence *in silico* and making a second copy of this sequence at with specific levels of variation from the original. Sequencing reads (100bp) were then generated *in silico* at a depth of one read every five base pairs. Reads were run through the assembly pipeline to assess whether the original DNA sequences were reconstructed from the read data, or if the reads formed a chimeric sequence. This simulation was repeated 250 times. The current assembly approach reliably reproduced single copy exons at all levels of variation and was able to re-assemble the individual paralogs where there was >6% divergence between original paralogous sequences, with inconsistent results at <5% divergence (**Fig. S4B**).

2.4. Contig merging and filtering

Sequencing reads were aligned to the assembled contigs using NextGenMap (v0.5.5; -R 40) (*11*). This alignment step allows for the recruitment of new reads to the contig that may have not been previously identified by blastn due to high degrees of variance relative to the reference blast database, large indels, or low amounts of overlapping sequence with the target region. This allows the contig to be elongated to include more of the flanking regions surrounding each target element. Reads were removed from the alignment to the contig if there were >3 mismatches with the exception of indels. To refine and extend the boundaries of the original contig, a second *de novo* assembly by CAP3 (-o 20 -p 85) was performed using the aligned reads.

Multiple contigs were present in around 65% of target regions after CAP3 assembly. To remove misidentified contigs, we used blastn to compare each contig to the reference genome, removing contigs whose top hit did not match the original bin from which the reads were assembled. To correct for potential assembly artifacts, the multiple contigs that represent each target were compared to each other and the reference sequence in a multiple sequence alignment using Mafft v7.313 (*12*) (--maxiterate 1000 --localpair), adding contigs as fragments (--addfragments). Using the read support at each base in the alignment, we generated a consensus contig sequence for each target region. A mismatch between contigs was considered if the read support for the most common base was <80% of all bases present. Contigs were only merged if there were <3 mismatches, if the sequence identity between the contigs was >95%, or if the contigs did not overlap within the target region. After this refinement step, <2% of target regions were represented by multiple contigs.

#### 3. Analysis of assembled reference contigs

##### 3.1 Comparison of assembled contigs to reference genome

In comparing our reference exome sequence assembly to the published *N. coriiceps* reference genome, we found 99.8% average percent identity of exome sequence to the reference target (**Fig. S5**), that included 98.5% of exons in the *N. coriiceps* reference genome (**Fig. S5**). The small differences in sequence identity and putative CNVs between this exome and the genome assembly may reflect meaningful biological variation in our independently sampled *N. coriiceps* populations. Both the sequence composition of the assembled exome and the predicted copy number data closely match the whole genome sequence data, providing confidence in our assembly.

##### 3.2. Estimation of read coverage and depth of targeted regions

Coverage was estimated using BEDtools (2.23.0) (*6*). Reads were first aligned to the assembled contigs with NextGenMap (*11*). The coordinates of the read alignments were then lifted the corresponding position on the reference genome using information from a pairwise sequence alignment between the contig and the orthologous region on the reference genome. Pairwise alignments were performed using Biopython v1.70 (pairwise2; match = 5, mismatch = -4, gap_open = -15, gap_extend = -1). Alignments were converted to BAM files, sorted, and indexed using SAMtools v1.9 (*13*). Reads alignments were manually inspected in the Integrative Genome Viewer (IGV) to verify accurate read alignment (*14*). Coverage is defined as the percentage of targeted bases in the primary reference genome having at least one read. Depth was estimated using coverageBed (-d).

##### 3.3. Distribution of read coverage across the dataset

Most target regions had either 0% or 100% coverage (**Fig. S6**). Though the average depth is similar in the notothenioids compared to the outgroups, there is a wider distribution of depths in outgroup species (**Fig. S6**). Though global coverage is >85% in all species, gene classes associated with the immune system, cell adhesion proteins and extracellular matrix were enriched among regions with relatively poor coverage (<25% coverage in >75% of exons; **Table S10**). This suggests that these fast evolving gene classes are less likely to be highly represented in these datasets, and is similar to previous findings with cross-species targeted DNA enrichment[67].

##### 3.4. Recovery of population variation

To determine the ability of this approach to recover population variation, we looked for heterozygous SNPs within the targeted regions of the dataset. Sequencing reads were aligned to the reference contigs for each species using NextGenMap (*11*). SAM files were converted to BAM files using SAMtools v1.9 (*13*). Variants were called using SAMtools mpileup and BCFtools v1.9

(call -mv). We considered sites heterozygous in our small population samples if there is ≥2 reads showing the variant in at least 25% allele frequency within the sequencing reads.

#### 4. Identification of orthologs

##### 4.1. Treatment of exons with a predicted history of duplication

We assembled a single exon/CNE copy for the majority of targets that were directly compared between species. However, less than 3% of targeted regions on average had assembled >1 contig per target region after assembly. For duplicated regions, both exon versions were ignored for that particular species in downstream analyses involving multiple groups, unless those analyses ask specific questions involving copy number.

##### 4.2. Identification of orthologous sequences

For ortholog pairing, the contigs generated from the same reference target region were aligned using Mafft v7.313(--op 10 --ep 10) (*12*), with a maximum likelihood tree topology estimated IQTree (*15*). Gene trees were reconciled with the species tree using Notung-2.9 (*16*) (--reconcile --rearrange --silent --threshold 90% --treeoutput nhx) to infer patterns of duplication and loss. The total number of duplication and loss events inferred by Notung were then summed and compared to a null scenario where all copies are local duplicates. If Notung inferred fewer gain/loss events, the duplicate exons were paired based on the reconciled gene tree.

##### 4.2. Simulation of ortholog identification approach

To estimate the ability of this approach to parse copy number variation into orthologous groups, we performed a series of simulations (**Fig. S7A**) using a random DNA sequence generated *in silico*. This ancestral DNA sequence was duplicated, and mutations were added at defined levels to distinguish each paralog. Both paralogs were then evolved according to a specified phylogeny, with variation added to each paralog in increments at each branch point. We varied copy number, length of contig sequence and also simulated local losses of an individual paralog within downstream lineages. Each simulation was repeated 250 times. These results suggest that as long as there is ≥4-6% variation between paralogous sequences, the approach can properly pair orthologous sequences (**Fig. S7B-D**). This coincides with the thresholds at which our pipeline can distinguish copy number variants during contig assembly (see above), meaning there will not be paralog sequences with <4% divergence in the dataset.

#### 5. Processing of reference contigs for comparative genomics

##### 5.1. Multiple sequence alignment

All paired orthologous sequences were aligned using Mafft v7.313 (--maxiterate 1000 --localpair --op 10 --ep 10). For coding regions that had out-of-frame or frameshift-causing indels, these alignments were then refined into codon alignments using the frameshift-aware multiple sequence aligner MACSE v2.03 (-prog alignSequences -seq -seq_lr -fs_lr 10 -stop_lr 15) (*17*). The multiple sequence alignment was pruned using GUIDANCE v2.02 to mask residues with score <0.6 (--bootstrap 25 ,--mafft, --maxiterate 100, --localpair --op 10 --ep 10 ) (*18*).

5.2. Reconstruction of gene sequences from exon data

Single-copy coding exons with orthology to *Gasterosteus aculeatus* were concatenated into gene sequences using the annotations of the *G. aculeatus* genome. The exons for each gene were spliced together in the same order in which they appear in the genome on the strand containing the gene. Transcript isoforms were merged into a non-redundant gene sequence containing all possible exons. A total of 18,600 gene sequences with orthology to *G. aculeatus* were reconstructed for each species.

#### 6. Comparative evolutionary analysis of genetic data

6.1. Notothenioid phylogeny

We used two approaches to infer a phylogeny for notothenioids. For both analyses, only single copy genes with >85% coverage in all species were included, resulting in a dataset of 11,627 genes. First, we individually partitioned each gene by codon position and used ModelFinder as implemented in IQTree v.1.6.3 (*19*) to estimate the optimal partitioning scheme and molecular evolution model. Genes were concatenated all a maximul likelihood tree was inferred using IQTree v.1.6.3 (*15*). To assess support for the phylogenetic relationships, we performed 1,000 ultra-fast bootstrap replicates (*20*).

In order to account for the effects of incomplete lineage sorting and known issues with concatenation for phylogenetic inference (*21*, *22*), we also inferred a species tree. Full species tree inference is not computationally feasible with large genomic datasets, so we relied on the summary species tree approach in ASTRAL v.5.6.2 (*23*). We first used IQTree v.1.6.3 to infer the maximum likelihood tree for each gene, apply partitioning schemes and molecular evolution models as described above. We then used ASTRAL to summarize the distribution of gene trees and estimate the species tree. To assess support for the species tree topology, we estimated local posterior probabilities for each quadpartition in the tree (*24*).

6.2. Quantification of lineage diversification dynamics

To test for changes in lineage diversification rates across the temporal history notothenioids, we used a Bayesian analysis of macroevolutionary mixtures implemented in BAMM v2.5 with a previously published time-tree that sampled all major lineages and 87 of ~120 species of notothenioids (*25*). Priors were were defined using the function setBAMMpriors contained in the R package *BAMMtools* v2.1.0 (*26*), and missing species were accounted for based on currently described species. BAMM runs were assessed for target sampling of the posterior distribution (Effective sample size >200) and results visualized using functions available from the R-package *BAMMtools* (*26*).

6.3. dS and dN estimates of substitution rate

dN, dS and dN/dS were calculated pairwise between each species and the outgroup *Percina caprodes*. This was performed for each reconstructed gene of at least 2,000 bp in both species using cal_dn_ds in the Biopython v1.70 codonseq module (method="NG86"). The values of dN, dS and dN/dS were then averaged across all genes for each species.

6.4. Molecular clock models of substitution rate

The substitution rate was estimated on the reconstructed gene sequences using the random local clock model as implemented in BEAST v2.4.8 (codon partitioned, bModelTest, chain length = 30M) (*27–29*). To simplify comparisons between gene trees, and as support for the relationships between the included species is high, we fixed the starting tree topology for each gene tree to match our ASTRAL-inferred species tree (**Fig. S8**). Only gene trees with ESS values ≥200 for all parameters were selected for downstream analysis (1,062 total). MRCA age priors were calibrated based on previous age estimates (*25*, *30*): *Psuedaphritis* + Eleginopsioidea 63.0my (52.6-73.4), Bovichtidae + all notothenioids 85.7my (69.5-102.6), *Harpagifer*-*Pogonophryne* 10.2my (7.7-13.0), *Bathydraco*-*Chaenocephalus* 11.1my (9.4-13.3), *Notothenia* 17.7my (15.2-20.5), Cryonotothenioidea 21.6my (18.6-23.9), Eleginopsiodea 45.9my (37.2-53.2). The maximum clade credibility was constructed for each gene tree using TreeAnnotator.

6.5. Detection of diversifying selection

Positive selection was calculated using the adaptive branch-site random effects likelihood (aBSREL) implemented in HyPhy v2.3.9 (*31*, *32*). Single copy exon alignments were concatenated into genes as input based on the gene order in the stickleback genome. The species tree was used for all comparisons (**Fig. S8**). Accelerated sequence evolution was assessed using phyloP (*33*) as implemented in PHAST v1.4 (*34*) (--method LRT --no-prune --features --mode ACC). The tree model for phyloP was derived separately for CNE and coding gene comparisons using phyloFit and the species (**Fig. S8**). Tree models for protein coding regions were based on 3,381 exons ≥1,000bp with ≥85% coverage in all species. CNEs tree models based on 2,912 elements ≥250bp with ≥85% coverage in all species.

6.6. Ontology enrichment

As there are no gene ontology (GO) terms relating to *Notothenia coriiceps*, we generated a custom gene ontology database based on the combined GO data from multiple species. GO data was mined from chicken, mouse, rat, human, stickleback, medaka and zebrafish in Ensembl BioMart (downloaded December 2016) (*35*). All Ensembl gene IDs were converted to their Ensembl stickleback ortholog and a final merged GO list was created by combining stickleback orthologs for each species. As many of the evolved phenotypes in notothenioids are comparable to human pathologies, we further utilized Human Phenotype Ontology databases to characterize genetic trends within the fish dataset (*36*). This HPO database (downloaded April 2018) was then converted from human to stickleback orthologs using Ensembl BioMart.

Ontology enrichment was performed using Fisher's Exact Test (SciPy v0.18.1; fisher_exact). We also assessed patterns of cumulative polygenic enrichment within ontologies using the SUMSTAT approach as implemented in Daub et al (*37*). This approach normalizes the distribution of log-likelihood ratio test values (ΔlnL) output from phyloP and HyPhy by taking the fourth root (ΔlnL4). The ΔlnL4 score is then summed for all genes within an ontology and an enrichment p-value is estimated from the empirical sum(ΔlnL4) score through bootstrap resampling (1,500 replicates). For all enrichment analyses, p-values were corrected using FDR (python module statsmodels v0.6.1; fdrcorrection0).

#### 7. Experimental analysis of bone density

##### 7.1. Zebrafish husbandry and genetic lines

Zebrafish wild-type and mutant lines used were housed and maintained as previously described and in accordance with Boston Children's Hospital IACUC regulations (*38*). The *col1a1a* (dmh14) and *col2a1* (dmh15) mutants were derived from a forward genetic screen (*39*).

##### 7.2. Zebrafish genome editing

The gRNA site GGTCAGAGTTTGGGTCAGGTCGG in exon 1 of the zebrafish *trip11* gene was targeted. This site is 33 bp downstream of the ATG start codon of *trip11*. gRNA sequences were cloned in the BsaI site of the DR274 plasmid and *in vitro* transcribed with T7 RNAmax kit (ThermoFisher) Cas9 mRNA was obtained from SystemBio (CAS500A-1). Injections of fertilized zebrafish eggs were done with 50 ng/ul gRNA and 150 ng/ul Cas9 mRNA. Genotyping was performed using 5'-CCCTGGTCGGTGATTTAGGGTTAG-3' as forward primer and 5' CACCTCCCACTTCCTCGGCGCTTTCCAGCAGAATATCTTTGGTAAAATTAGAG-3' as reverse primer. PCRs using this primer pair yield a 178 bp wildtype band. Fish were identified with a 47 bp deletion spanning the gRNA target site and yielding a 131 bp genotyping band. The deleted sequence is 5' -CTGGGTCAGAGTTTGGGTCAGGTCGGGGGAAGCTTGTCTTCATTTAC-3' and generates a frameshift at the 12th amino acid residue of *trip11*.

##### 7.3. Analysis of skeletal density through computed tomography (CT)

Adult notothenioid specimens were loaned from the Yale Peabody Museum (**Table S11**) and scanned using the Siemens Biograph at Boston Children's Hospital Department of Nuclear Medicine and Molecular Imaging. Scan data was processed in Siemens PETsyngo VG60A software and analyzed in Amira (v 6.0.0 ; FEI Inc.).

Zebrafish were euthanized in 22% MS-222, fixed in 3.7% formaldehyde overnight, and rinsed in PBS. The fishes were embedded in 1% agarose to reduce movement during imaging. The fish skulls were scanned as in ref. 66, using a Skyscan 1173 (Bruker), 240-degree scan with 0.2 rotational step. X-ray source voltage set to 70 kV and current set to 80 μA. Exposure time was 1500 ms. Resolution of scan was 7.14 microns per pixel. Volume renderings were reconstructed as maximum intensity projections in Amira software. Skeletal density was estimated based on the average pixel intensity measured from the maximum intensity projection of the skull, vertebrae and operculum in ImageJ v1.51s (https://imagej.nih.gov/ij/). A total of n=6 *trip11*$^{+/-}$ and n=10 *trip11*$^{-/-}$ fish were scanned and quantified. The calvaria, operculum and vertebrae were measured from each individual.

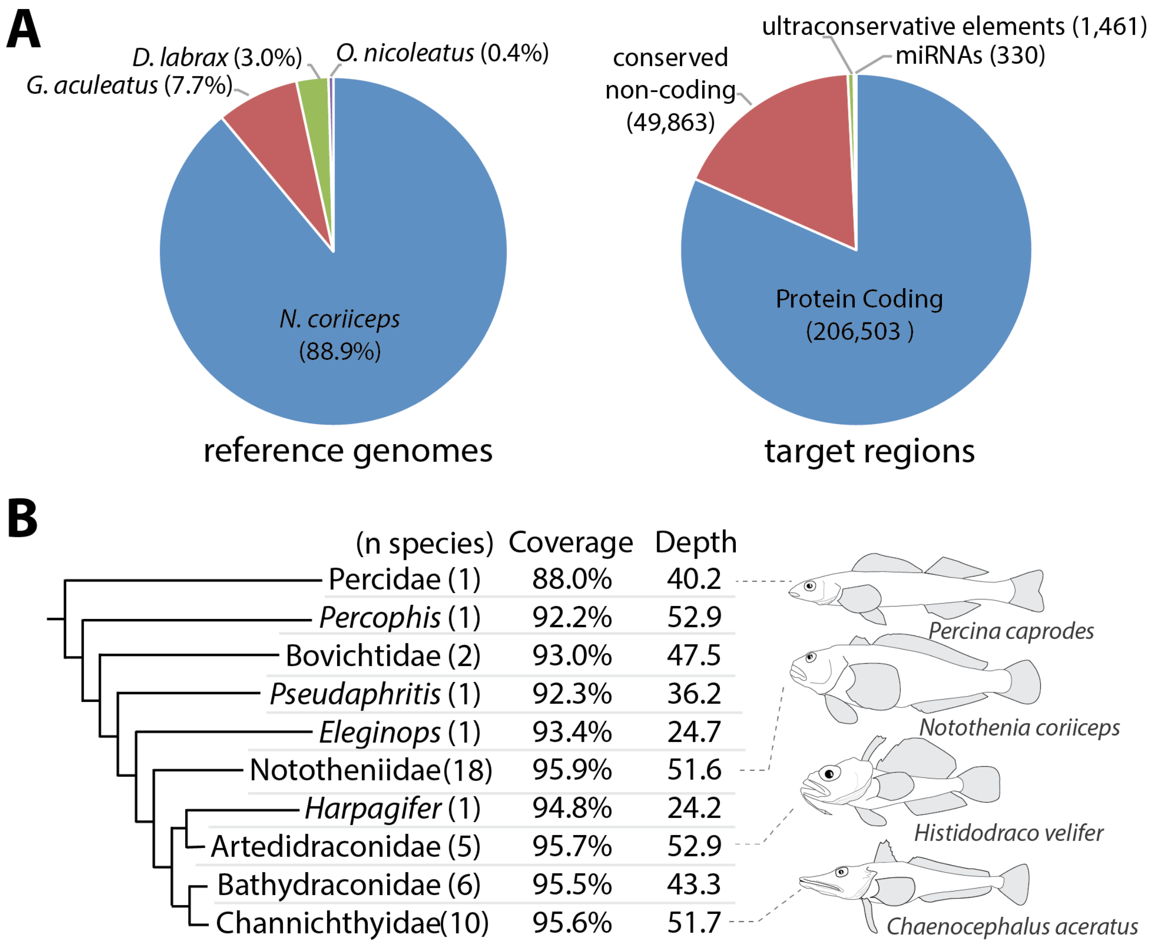


**Fig. S1. Evolutionary history of notothenioids revealed through targeted sequence enrichment**. (**A**) Designed 'phylo-capture' array demarcating regions targeted for cross-species sequence enrichment; total number of elements in parentheses. (**B**) Average coverage and sequencing read depth of the targeted regions for each major group within the notothenioids and close outgroups. The number of species within each group is shown in parenthesis.

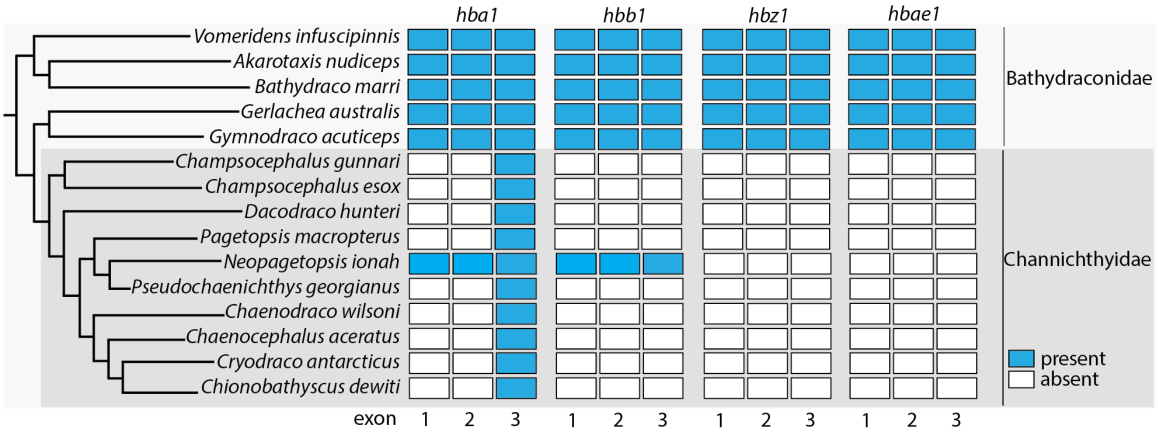


**Fig. S2. Sensitivity of targeted capture method in confirming and discovering variation in hemoglobin within Channichthyidae**. Presence and lack of detection of individual hemoglobin exons in Channichthyidae and Bathydraconidae. Genomic analysis was sensitive in detecting loss of hemoglobins *hba1* and *hbb2* in the icefish clade as previously identified (*40*). The analysis was able to identify patterns of loss within two new hemoglobins *hbz1* and *hbae1* as well. Presence defined as > 25% coverage and > 5 sequencing reads per exon.

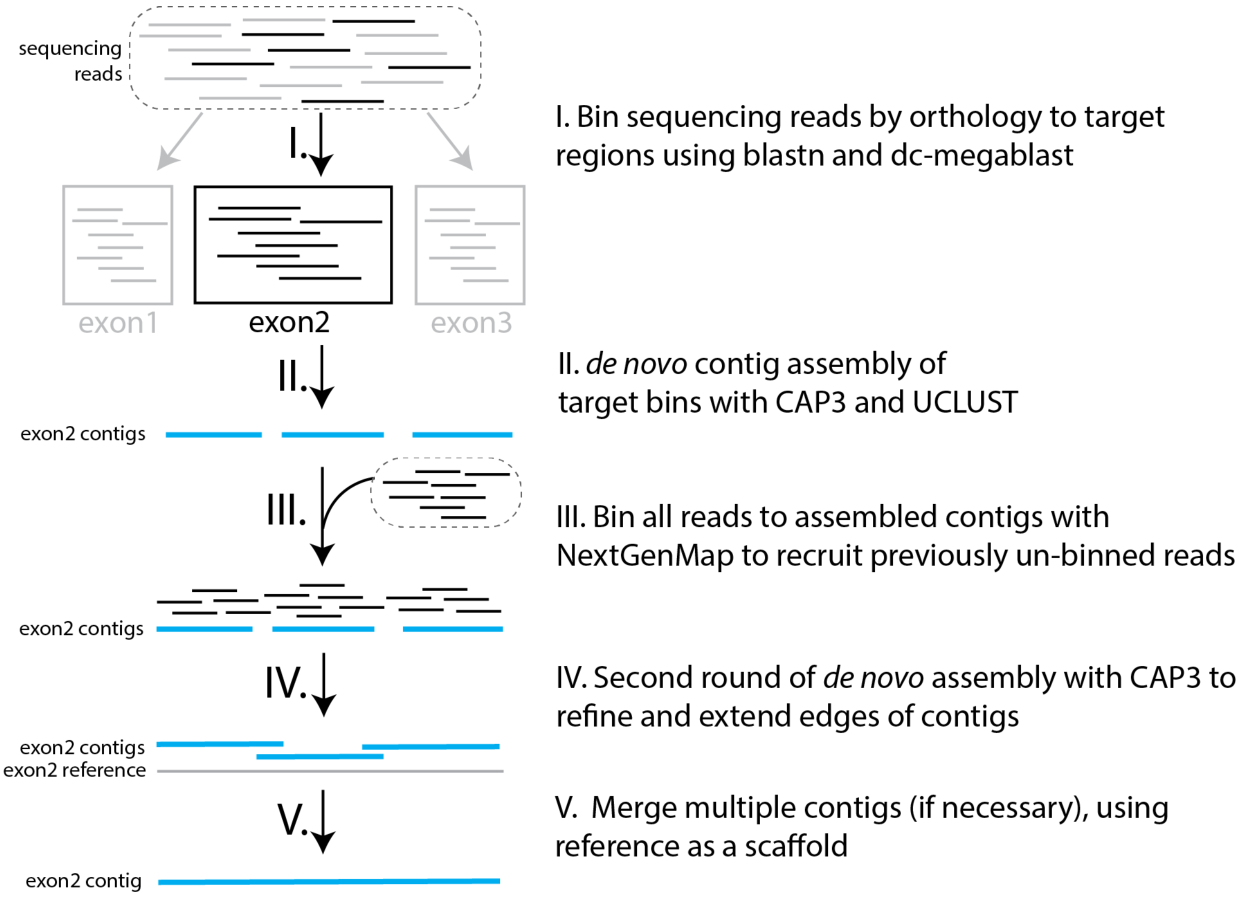


**Fig. S3. Overview of contig assembly**

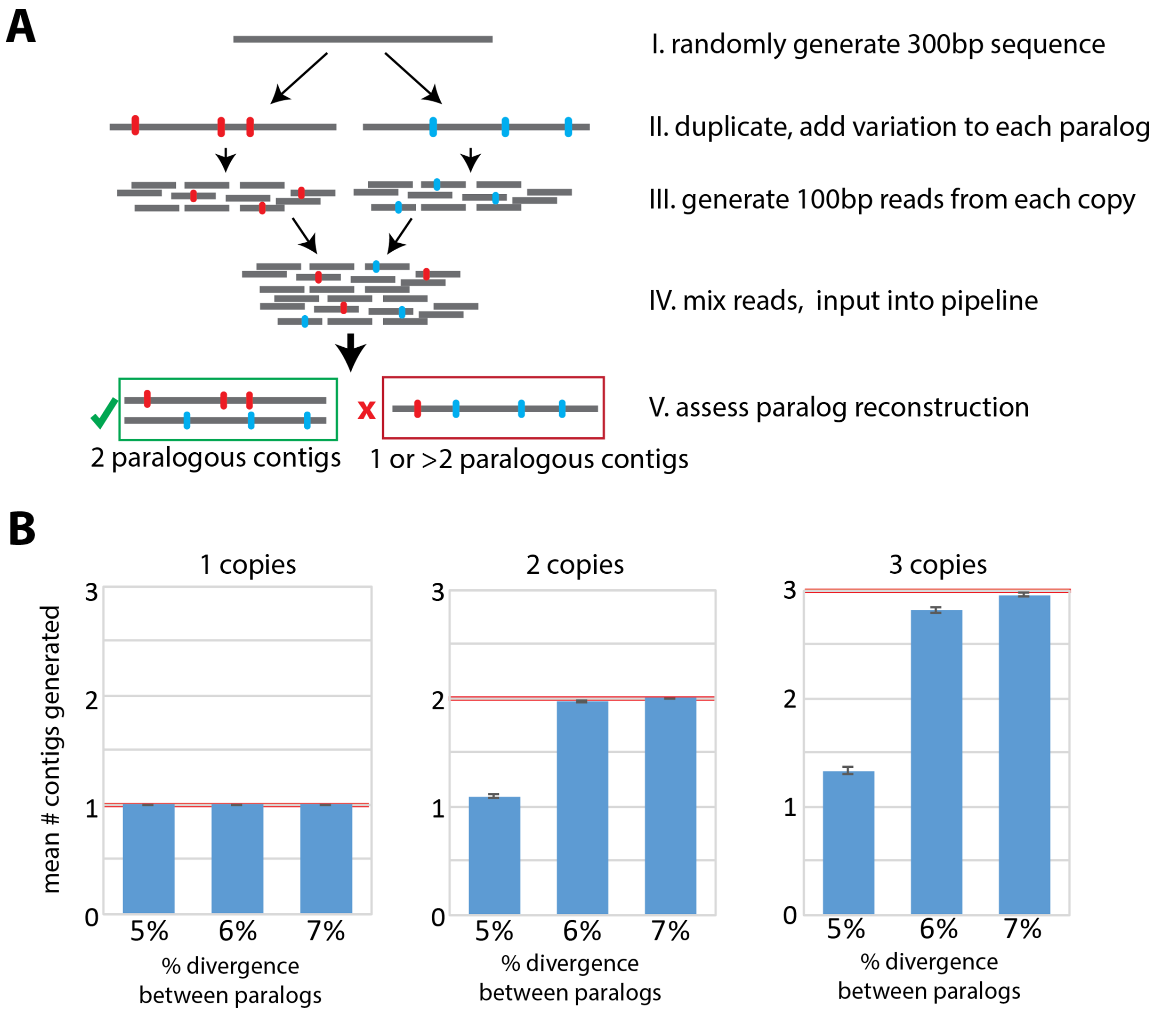


**Fig. S4. Detection of copy number variants through contig assembly**. **(A)** Simulation of the limits of the contig assembly pipeline to reconstruct duplicate sequences from mixed read populations. (**B**) The mean number of contigs generated by the pipeline from 1, 2 and 3 input contigs put into the simulation. Data from 250 simulations of random 300-bp DNA fragments. Red line indicates input number of contigs. Error bars represent ± 1 sem.

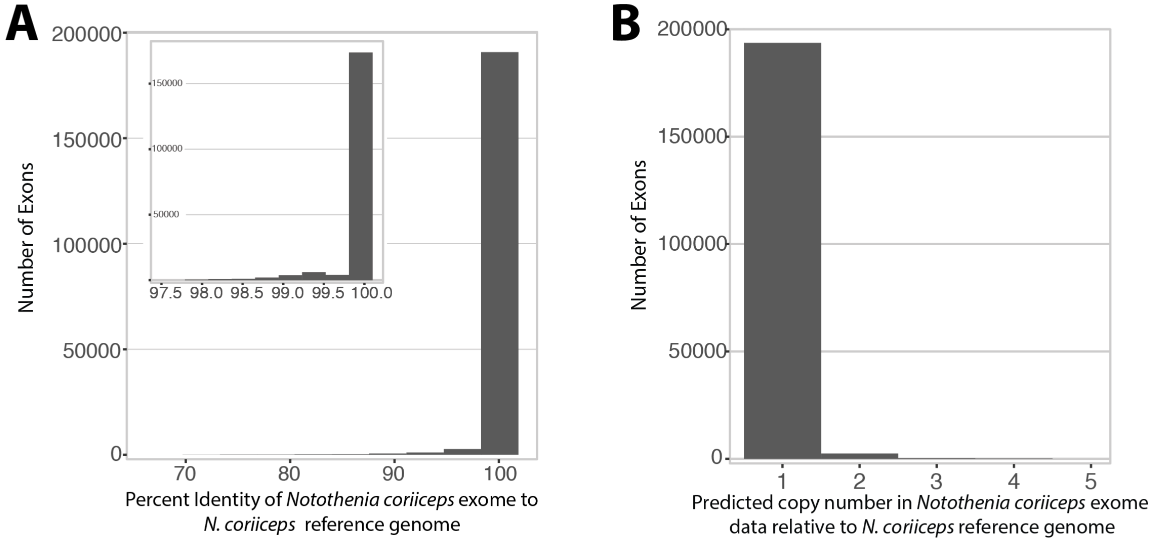


**Fig. S5. Comparison of reconstructed *Notothenia coriiceps* exome to published *N. coriiceps* genome.** (**A**) Histogram of percent identity of the reference target regions relative to the corresponding region on the previously assembled *N. coriiceps* genome (*1*). (**B**) Histogram of predicted copy number for an exon in the exome reconstruction compared with the reference genome.

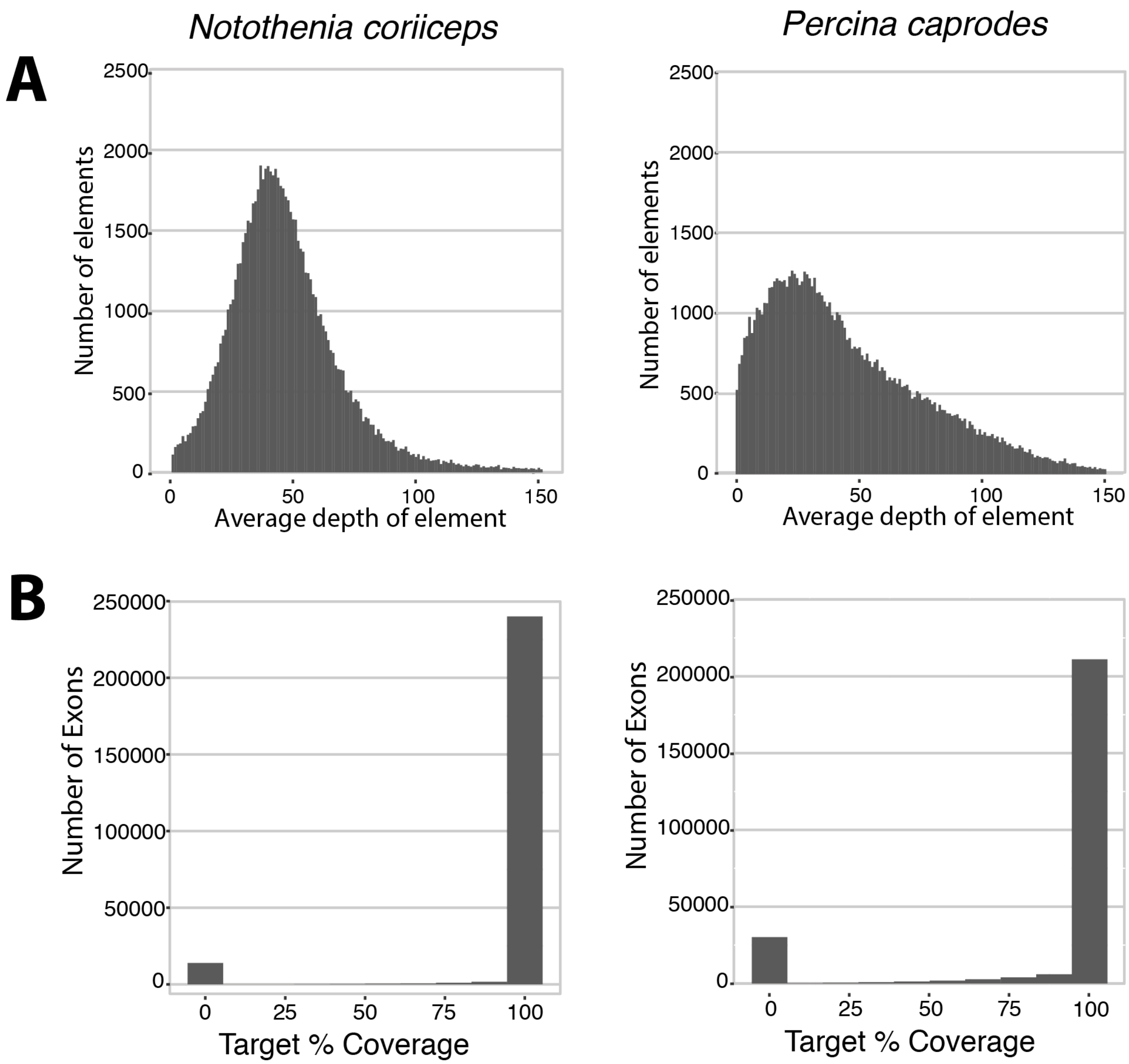


**Fig. S6. Distribution of coverage as a function of evolutionary distance from hybridization baits**. (**A**) Histogram of average depth per targeted region (exon, CNE, etc.), showing flattened distribution of read depth in *P. caprodes* suggesting that certain target regions in distantly related groups hybridize less efficiently. (**B**) Percent of bases covered with at least one read in both species, showing the majority of target regions were either highly covered or not covered at all.

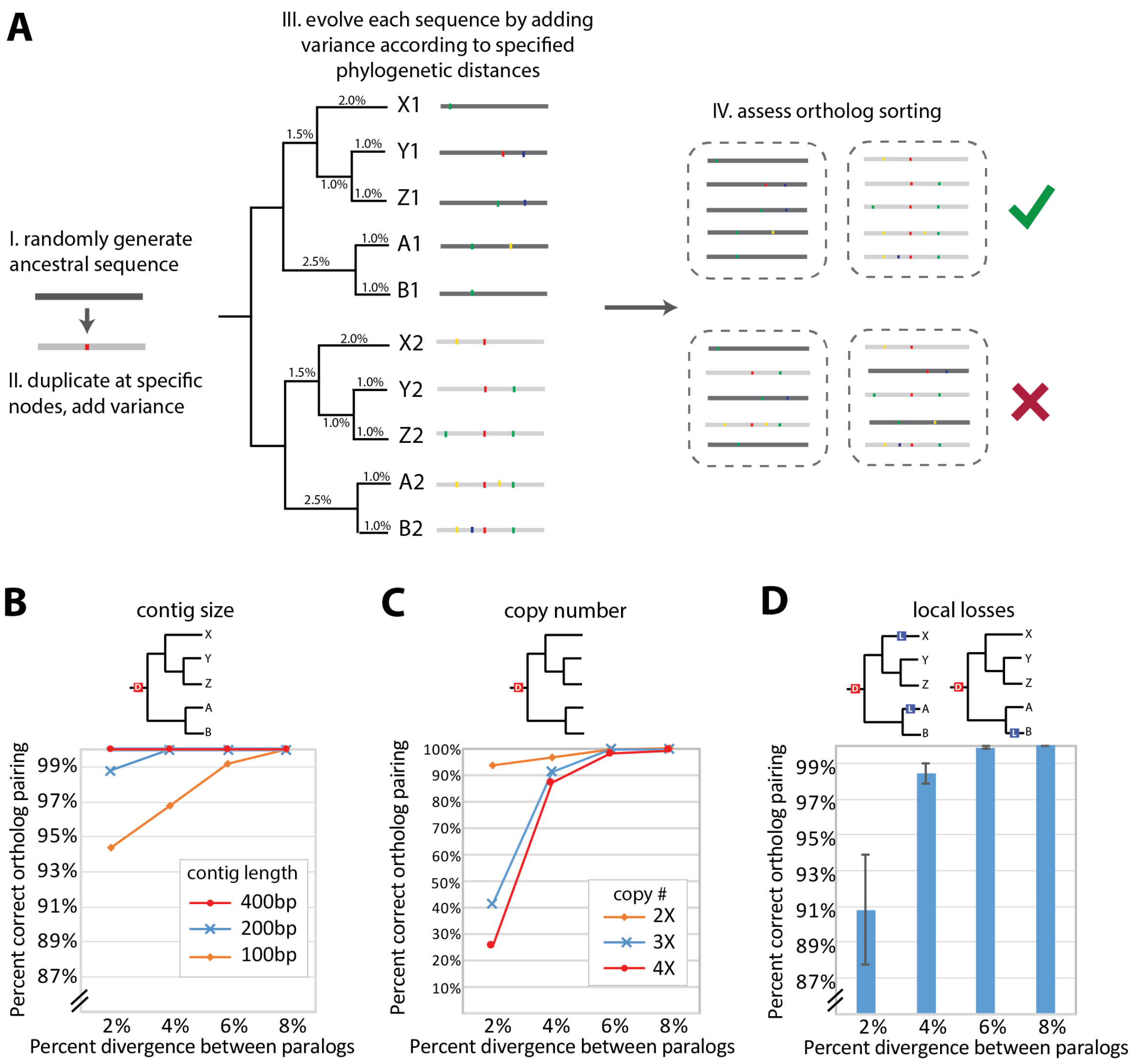


**Fig. S7. Assessing robustness of pairing of duplicated elements for comparative genomics.** (**A**) Simulation to test the ability of this approach to properly identify orthologs among ancestrally duplicated sequences. As part of the pipeline for annotation, when there is more than one element per target region in the reference genome in at least two species, a gene tree is constructed with IQTree and reconciled with the species tree using Notung. Notung infers patterns of gain and loss throughout the phylogeny. Simulation was run 250 times. **B-D**) Effect of element length (**B**), copy number (**C**) and local losses within the phylogeny (**D**) on correct matching of duplicates relative to the percent divergence of the duplicated elements. Error bars represent ± 1 sem.

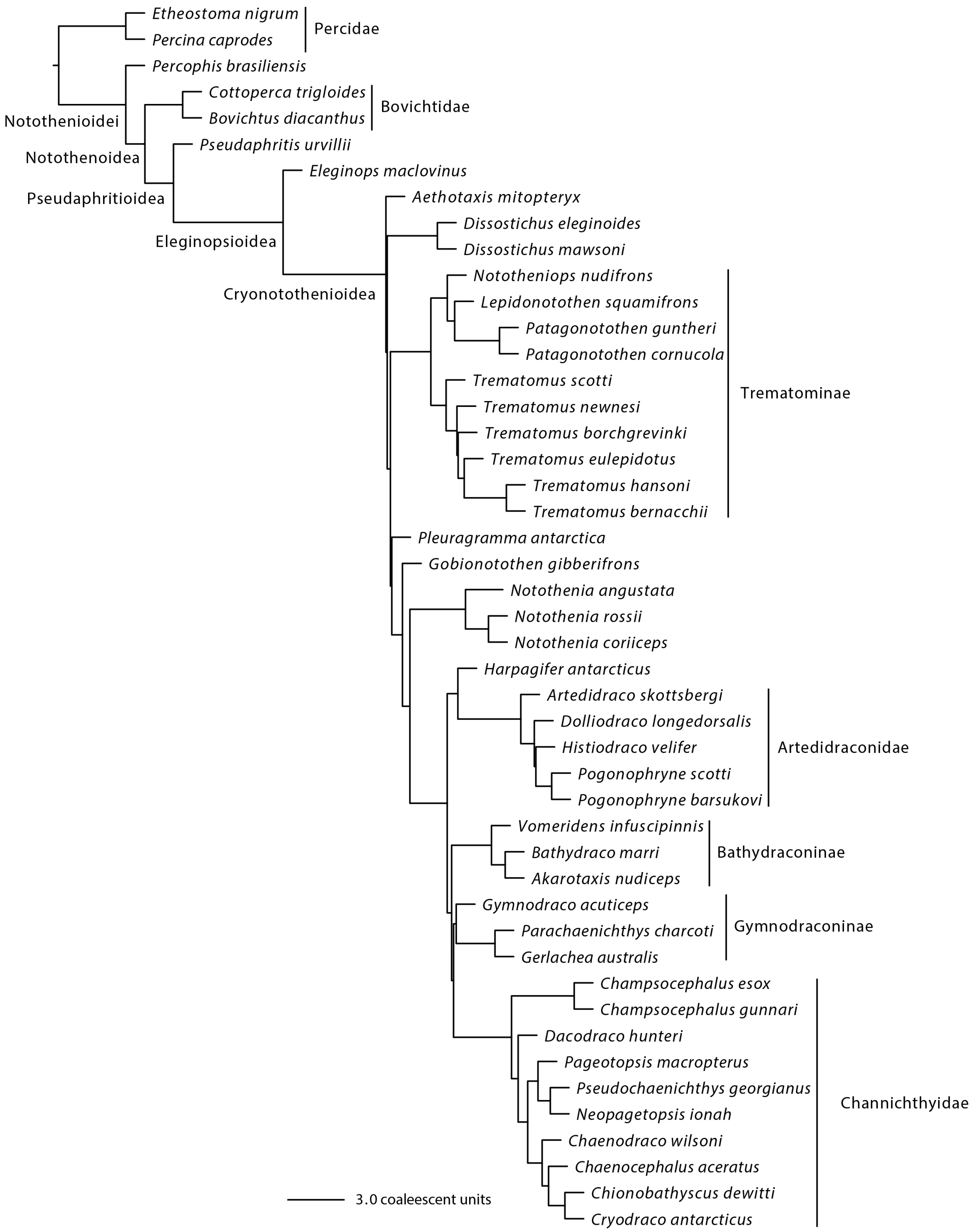


**Fig. S8. Notothenioid species tree.** Phylogenetic relationships of the sequenced species inferred as a species tree using ASTRAL and 11,627 gene trees. All nodes were supported with a quadpartition posterior probability of 1.00.

Table S1. Coverage at various read depth thresholds

| | Coverage at depth | | | |
|---|---|---|---|---|
| **Species** | **>1X** | **>2X** | **>4X** | **>10X** |
| *Percina caprodes* | 88.0% | 87.4% | 84.5% | 73.4% |
| *Percophis brasiliensis* | 92.2% | 92.0% | 90.4% | 83.7% |
| *Bovichtus diacanthus* | 92.2% | 91.9% | 90.6% | 82.8% |
| *Cottoperca trigloides* | 93.9% | 92.4% | 88.6% | 78.3% |
| *Pseudaphritis urvillii* | 92.3% | 91.8% | 89.4% | 79.1% |
| *Eleginops maclovinus* | 93.4% | 93.0% | 90.9% | 78.2% |
| *Pleurgramma antarctica* | 95.9% | 95.8% | 95.3% | 92.7% |
| *Aethotaxis mitopteryx* | 96.4% | 96.4% | 96.1% | 95.2% |
| *Dissostichus mawsoni* | 95.6% | 95.5% | 94.7% | 88.4% |
| *Dissostichus eleginoides* | 95.9% | 95.8% | 95.4% | 93.0% |
| *Nototheniops nudifrons* | 95.9% | 95.8% | 95.4% | 93.3% |
| *Lepidonotothen squamifrons* | 95.9% | 95.9% | 95.5% | 93.6% |
| *Patagonotothen squamiceps* | 95.9% | 95.9% | 95.5% | 93.6% |
| *Patagonotothen guntheri* | 95.9% | 95.8% | 95.4% | 93.3% |
| *Trematomus scotti* | 95.7% | 95.6% | 95.1% | 92.2% |
| *Trematomus newnesi* | 95.8% | 95.7% | 95.2% | 92.3% |
| *Trematomus eulepidotus* | 95.9% | 95.9% | 95.5% | 93.9% |
| *Trematomus bernacchii* | 95.8% | 95.6% | 95.1% | 91.6% |
| *Trematomus borchgrevinki* | 95.7% | 95.6% | 95.1% | 90.8% |
| *Trematomus hansoni* | 95.6% | 95.5% | 95.1% | 92.6% |
| *Gobionotothen gibberifrons* | 96.0% | 95.9% | 95.4% | 93.1% |
| *Notothenia coriiceps* | 96.0% | 95.9% | 95.5% | 93.3% |
| *Notothenia rossii* | 95.7% | 95.6% | 95.1% | 91.8% |
| *Paranotothenia angustata* | 96.0% | 95.9% | 95.4% | 92.2% |
| *Harpagifer antarcticus* | 94.8% | 94.6% | 93.1% | 81.9% |
| *Artedidraco skottsbergi* | 95.3% | 95.1% | 94.2% | 88.0% |
| *Histiodraco velifer* | 95.8% | 95.7% | 95.3% | 93.3% |
| *Dolliodraco longedorsalis* | 95.8% | 95.7% | 95.4% | 94.1% |
| *Pogonophryne scotti* | 95.7% | 95.6% | 95.2% | 92.8% |
| *Pogonophryne barsukovi* | 95.8% | 95.7% | 95.3% | 93.1% |
| *Gerlachea australis* | 95.6% | 95.4% | 94.5% | 88.9% |
| *Parachaenichthys charcoti* | 94.2% | 93.8% | 92.0% | 79.8% |
| *Vomeridens infuscipinnis* | 95.8% | 95.7% | 95.1% | 91.7% |
| *Akarotaxis nudiceps* | 95.8% | 95.7% | 95.1% | 91.9% |
| *Bathydraco marri* | 95.7% | 95.6% | 95.2% | 92.3% |
| *Gymnodraco acuticeps* | 96.2% | 96.1% | 95.8% | 94.4% |

| | | | | |
|---|---|---|---|---|
| *Dacodraco hunteri* | 95.4% | 95.4% | 94.9% | 92.3% |
| *Champsocephalus gunnari* | 95.7% | 95.5% | 94.9% | 92.0% |
| *Champsocephalus esox* | 95.4% | 95.3% | 94.7% | 91.5% |
| *Pageotopsis macropterus* | 95.4% | 95.3% | 94.7% | 91.6% |
| *Neopagetopsis ionah* | 95.5% | 95.4% | 94.8% | 91.3% |
| *Pseudochaenichthys georgianus* | 95.4% | 95.3% | 94.8% | 92.1% |
| *Chaenodraco wilsoni* | 95.5% | 95.4% | 95.0% | 93.0% |
| *Chaenocephalus aceratus* | 96.0% | 95.9% | 95.6% | 93.9% |
| *Cryodraco antarcticus* | 95.6% | 95.5% | 94.9% | 91.5% |
| *Chionobathyscus dewitti* | 95.2% | 94.8% | 93.7% | 88.6% |

### Table S2. Coverage of targeted regions in cross-species targeted sequence enrichment

| | total | | *N. coriiceps* genome | | non-*N.coriiceps* genome | | CDS | | CNE | |
|---|---|---|---|---|---|---|---|---|---|---|
| species | cov. | depth | cov. | depth | cov. | depth | cov. | depth | cov. | depth |
| *Percina caprodes* | 88.0% | 40.2 | 87.6% | 34.9 | 87.4% | 60.3 | 86.6% | 34.4 | 94.8% | 64.7 |
| *Percophis brasiliensis* | 92.2% | 52.9 | 92.3% | 46.7 | 90.3% | 76.3 | 91.3% | 45.6 | 96.6% | 84.7 |
| *Bovichtus_ diacanthus* | 92.2% | 54.3 | 92.3% | 47.5 | 89.9% | 78.7 | 91.3% | 46.1 | 96.1% | 90.4 |
| *Cottoperca trigloides* | 93.9% | 40.7 | 94.4% | 35.8 | 89.7% | 59.4 | 93.4% | 35.0 | 95.9% | 66.4 |
| *Pseudaphritis urvillii* | 92.3% | 36.2 | 92.5% | 32.2 | 88.7% | 51.4 | 91.5% | 31.4 | 96.1% | 57.5 |
| *Eleginops maclovinus* | 93.4% | 24.7 | 94.3% | 23.4 | 85.4% | 28.9 | 92.9% | 23.0 | 95.4% | 32.2 |
| *Aethotaxis mitopteryx* | 95.9% | 48.5 | 97.2% | 48.8 | 86.3% | 43.7 | 95.8% | 48.8 | 96.4% | 47.1 |
| *Dissostichus mawsoni* | 96.4% | 89.1 | 97.7% | 89.6 | 87.9% | 81.3 | 96.4% | 89.3 | 96.7% | 87.8 |
| *Dissostichus eleginoides* | 95.6% | 29.9 | 97.0% | 30.3 | 85.4% | 26.9 | 95.5% | 30.3 | 96.0% | 27.8 |
| *Nototheniops nudifrons* | 95.9% | 50.1 | 97.2% | 50.2 | 86.5% | 46.6 | 95.8% | 50.2 | 96.4% | 49.7 |
| *Lepidonotothen squamifrons* | 95.9% | 52.1 | 97.2% | 52.1 | 86.5% | 48.9 | 95.8% | 52.0 | 96.4% | 52.6 |
| *Patagonotothen squamiceps* | 95.9% | 53.0 | 97.2% | 53.2 | 86.9% | 49.6 | 95.9% | 53.1 | 96.4% | 52.6 |
| *Patagonotothen guntheri* | 95.9% | 54.5 | 97.2% | 54.8 | 86.9% | 50.5 | 95.8% | 55.1 | 96.4% | 51.8 |
| *Trematomus scotti* | 95.9% | 54.6 | 97.2% | 54.7 | 86.6% | 51.0 | 95.8% | 54.8 | 96.4% | 53.6 |
| *Trematomus newnesi* | 95.7% | 44.4 | 97.0% | 44.6 | 86.1% | 41.2 | 95.6% | 44.5 | 96.2% | 43.9 |
| *Trematomus eulepidotus* | 95.8% | 47.1 | 97.1% | 47.5 | 86.1% | 42.5 | 95.7% | 47.5 | 96.3% | 45.4 |
| *Trematomus bernacchii* | 95.9% | 54.6 | 97.2% | 55.4 | 86.5% | 47.8 | 95.8% | 55.5 | 96.4% | 50.3 |
| *Trematomus borchgrevinki* | 95.8% | 44.7 | 97.1% | 44.8 | 86.1% | 41.8 | 95.6% | 44.8 | 96.3% | 43.9 |
| *Trematomus hansoni* | 95.7% | 41.6 | 97.1% | 42.1 | 85.9% | 37.9 | 95.6% | 42.2 | 96.2% | 38.9 |
| *Pleuragramma antarctica* | 95.6% | 61.3 | 96.9% | 61.1 | 86.4% | 58.2 | 95.5% | 60.9 | 96.1% | 62.9 |
| *Gobionotothen gibberifrons* | 96.0% | 51.0 | 97.3% | 51.3 | 86.3% | 46.5 | 95.8% | 51.2 | 96.5% | 49.9 |
| *Notothenia coriiceps* | 96.0% | 54.3 | 97.5% | 54.7 | 85.0% | 49.3 | 95.8% | 54.6 | 96.8% | 53.0 |
| *Notothenia rossii* | 95.7% | 54.9 | 97.2% | 57.0 | 84.5% | 43.5 | 95.5% | 57.1 | 96.4% | 44.8 |
| *Paranotothenia angustata* | 96.0% | 43.1 | 97.4% | 44.0 | 85.6% | 36.9 | 95.9% | 43.7 | 96.6% | 40.6 |
| *Harpagifer antarcticus* | 94.8% | 24.2 | 96.4% | 24.5 | 83.3% | 21.4 | 94.7% | 24.4 | 95.4% | 23.1 |
| *Artedidraco skottsbergi* | 95.3% | 32.1 | 96.8% | 32.4 | 84.5% | 29.2 | 95.2% | 32.3 | 96.0% | 30.9 |
| *Histiodraco velifer* | 95.8% | 55.3 | 97.2% | 56.0 | 85.8% | 49.1 | 95.7% | 55.9 | 96.5% | 52.7 |
| *Dolliodraco longedorsalis* | 95.8% | 68.2 | 97.2% | 69.2 | 85.6% | 59.4 | 95.6% | 69.1 | 96.5% | 64.0 |
| *Pogonophryne scotti* | 95.7% | 52.6 | 97.1% | 53.1 | 85.6% | 47.0 | 95.6% | 53.0 | 96.5% | 50.5 |
| *Pogonophryne barsukovi* | 95.8% | 56.5 | 97.2% | 57.2 | 85.8% | 50.5 | 95.7% | 57.1 | 96.5% | 54.0 |
| *Gerlachea australis* | 95.6% | 35.2 | 97.0% | 35.4 | 84.9% | 32.4 | 95.4% | 35.3 | 96.2% | 34.6 |
| *Parachaenichthys charcoti* | 94.2% | 27.8 | 95.8% | 28.5 | 82.0% | 23.6 | 94.1% | 28.5 | 94.6% | 24.4 |
| *Vomeridens infuscipinnis* | 95.9% | 63.2 | 97.3% | 64.6 | 85.8% | 53.9 | 95.8% | 64.7 | 96.5% | 56.6 |
| *Akarotaxis nudiceps* | 95.8% | 44.2 | 97.2% | 44.7 | 85.5% | 39.8 | 95.6% | 44.6 | 96.4% | 42.4 |
| *Bathydraco marri* | 95.8% | 45.7 | 97.2% | 46.3 | 85.6% | 40.8 | 95.7% | 46.2 | 96.4% | 43.4 |

| | | | | | | | | | | |
|---|---|---|---|---|---|---|---|---|---|---|
| *Gymnodraco acuticeps* | 95.7% | 43.4 | 97.2% | 44.4 | 85.2% | 37.1 | 95.6% | 44.5 | 96.3% | 38.5 |
| *Champsocephalus gunnari* | 96.2% | 74.8 | 97.5% | 75.4 | 86.7% | 67.5 | 96.0% | 75.2 | 96.9% | 72.6 |
| *Champsocephalus esox* | 95.4% | 50.3 | 96.9% | 51.0 | 85.1% | 43.3 | 95.3% | 50.9 | 96.0% | 47.1 |
| *Dacodraco hunteri* | 95.7% | 47.8 | 97.1% | 48.1 | 85.3% | 43.3 | 95.6% | 47.9 | 96.3% | 47.1 |
| *Pageotopsis macropterus* | 95.4% | 46.0 | 96.8% | 46.7 | 84.9% | 40.4 | 95.3% | 46.5 | 96.0% | 43.6 |
| *Neopagetopsis ionah* | 95.4% | 48.0 | 96.8% | 48.6 | 85.0% | 42.5 | 95.3% | 48.5 | 96.0% | 45.7 |
| *Pseudochaenichthys georgianus* | 95.5% | 44.6 | 96.9% | 44.8 | 85.3% | 41.5 | 95.4% | 44.5 | 96.1% | 45.3 |
| *Chaenodraco wilsoni* | 95.4% | 48.3 | 96.8% | 48.9 | 84.8% | 43.0 | 95.2% | 48.8 | 96.1% | 45.9 |
| *Chaenocephalus aceratus* | 95.5% | 53.6 | 96.9% | 54.3 | 85.1% | 47.3 | 95.4% | 54.2 | 96.2% | 50.8 |
| *Cryodraco antarcticus* | 96.0% | 62.3 | 97.4% | 63.2 | 86.3% | 54.8 | 95.9% | 63.1 | 96.6% | 58.4 |
| *Chionobathyscus dewitti* | 95.6% | 41.7 | 97.0% | 42.3 | 85.3% | 36.6 | 95.5% | 42.3 | 96.1% | 38.6 |

### Table S3. Recovery of heterozygous and fixed variation

| Species | # Het SNPs | Het SNP/ base | Percent of targets with Het | # Fixed species-specific SNPs | Fixed SNP/ base | Percent of targets with fixed, species-specific SNP |
|---|---|---|---|---|---|---|
| *Percina caprodes* | 48,777 | 0.0014 | 15.7% | 300,691 | 0.0087 | 52.9% |
| *Percophis brasiliensis* | 69,209 | 0.0019 | 19.7% | 698,153 | 0.0187 | 77.4% |
| *Bovichtus diacanthus* | 36,886 | 0.0010 | 11.6% | 379,258 | 0.0096 | 54.1% |
| *Cottoperca trigloides* | 30,261 | 0.0008 | 8.6% | 428,614 | 0.0116 | 62.6% |
| *Pseudaphritis urvillii* | 31,342 | 0.0009 | 10.2% | 501,713 | 0.0137 | 69.9% |
| *Eleginops maclovinus* | 53,559 | 0.0014 | 16.1% | 599,422 | 0.0162 | 74.2% |
| *Pleuragramma antarctica* | 154,174 | 0.0039 | 31.7% | 223,284 | 0.0057 | 41.1% |
| *Aethotaxis mitopteryx* | 77,172 | 0.0020 | 19.7% | 69,895 | 0.0018 | 19.3% |
| *Dissostichus mawsoni* | 42,867 | 0.0011 | 12.1% | 40,827 | 0.0010 | 12.7% |
| *Dissostichus eleginoides* | 53,849 | 0.0014 | 15.6% | 44,805 | 0.0012 | 14.0% |
| *Nototheniops nudifrons* | 58,203 | 0.0015 | 15.6% | 41,278 | 0.0011 | 12.5% |
| *Lepidonotothen squamifrons* | 62,976 | 0.0016 | 17.3% | 26,827 | 0.0007 | 8.4% |
| *Patagonotothen squamiceps* | 44,838 | 0.0011 | 12.4% | 53,013 | 0.0014 | 16.0% |
| *Patagonotothen guntheri* | 74,580 | 0.0019 | 19.7% | 23,794 | 0.0006 | 7.5% |
| *Trematomus scotti* | 66,169 | 0.0017 | 17.7% | 25,783 | 0.0007 | 8.2% |
| *Trematomus newnesi* | 53,985 | 0.0014 | 14.9% | 39,385 | 0.0010 | 12.3% |
| *Trematomus eulepidotus* | 47,855 | 0.0012 | 12.2% | 25,282 | 0.0006 | 8.1% |
| *Trematomus bernacchii* | 58,849 | 0.0015 | 16.1% | 2,307 | 0.0001 | 0.4% |
| *Trematomus borchgrevinki* | 37,388 | 0.0010 | 10.3% | 33,673 | 0.0009 | 10.9% |
| *Trematomus hansoni* | 61,373 | 0.0016 | 17.1% | 2,041 | 0.0001 | 0.3% |
| *Gobionotothen gibberifrons* | 64,191 | 0.0016 | 16.3% | 111,759 | 0.0029 | 28.0% |
| *Notothenia coriiceps* | 60,749 | 0.0015 | 16.2% | 40,240 | 0.0010 | 12.6% |
| *Notothenia rossii* | 66,220 | 0.0017 | 17.8% | 45,190 | 0.0012 | 13.6% |
| *Paranotothenia angustata* | 55,302 | 0.0014 | 15.5% | 104,620 | 0.0027 | 27.7% |
| *Harpagifer antarcticus* | 43,842 | 0.0012 | 13.1% | 53,936 | 0.0014 | 16.4% |
| *Artedidraco skottsbergi* | 64,811 | 0.0017 | 18.5% | 11,638 | 0.0003 | 3.9% |
| *Histiodraco velifer* | 42,299 | 0.0011 | 12.1% | 8,164 | 0.0002 | 2.7% |
| *Dolliodraco longedorsalis* | 52,428 | 0.0013 | 15.0% | 5,526 | 0.0001 | 1.7% |
| *Pogonophryne scotti* | 45,650 | 0.0012 | 13.2% | 3,446 | 0.0001 | 1.0% |
| *Pogonophryne barsukovi* | 44,453 | 0.0011 | 12.8% | 3,061 | 0.0001 | 0.9% |
| *Gerlachea australis* | 73,269 | 0.0019 | 20.5% | 21,708 | 0.0006 | 7.0% |

| | | | | | | |
|---|---|---|---|---|---|---|
| *Parachaenichthys charcoti* | 30,350 | 0.0009 | 9.2% | 46,682 | 0.0013 | 14.4% |
| *Vomeridens infuscipinnis* | 41,820 | 0.0011 | 11.0% | 36,750 | 0.0009 | 11.8% |
| *Akarotaxis nudiceps* | 56,254 | 0.0014 | 15.6% | 12,176 | 0.0003 | 4.0% |
| *Bathydraco marri* | 48,433 | 0.0012 | 13.2% | 19,462 | 0.0005 | 6.4% |
| *Gymnodraco acuticeps* | 58,027 | 0.0015 | 16.5% | 41,434 | 0.0011 | 12.9% |
| *Dacodraco hunteri* | 52,739 | 0.0013 | 14.8% | 23,605 | 0.0006 | 7.8% |
| *Champsocephalus gunnari* | 37,061 | 0.0010 | 10.9% | 14,024 | 0.0004 | 4.5% |
| *Champsocephalus esox* | 35,548 | 0.0009 | 10.3% | 40,568 | 0.0010 | 12.7% |
| *Pageotopsis macropterus* | 41,887 | 0.0011 | 12.1% | 20,670 | 0.0005 | 6.9% |
| *Neopagetopsis ionah* | 58,368 | 0.0015 | 15.9% | 17,087 | 0.0004 | 5.6% |
| *Pseudochaenichthys georgianus* | 14,921 | 0.0004 | 3.7% | 29,936 | 0.0008 | 9.8% |
| *Chaenodraco wilsoni* | 108,985 | 0.0028 | 27.6% | 4,751 | 0.0001 | 1.4% |
| *Chaenocephalus aceratus* | 26,447 | 0.0007 | 7.1% | 18,353 | 0.0005 | 6.2% |
| *Cryodraco antarcticus* | 115,004 | 0.0029 | 26.9% | 2,531 | 0.0001 | 0.4% |
| *Chionobathyscus dewitti* | 44,558 | 0.0011 | 11.9% | 8,258 | 0.0002 | 2.7% |

### Table S4. Substitution rates in notothenioids and outgroups

| Branch | Substitution Rate | 95% HPD |
|---|---|---|
| *Percina caprodes* | 0.00078 | 0.0005-0.0011 |
| *Percophis brasiliensis* | 0.00077 | 0.0005-0.0011 |
| *Bovichtus diacanthus* | 0.00077 | 0.0005-0.0011 |
| *P. urvillii* + Eleginopsioidea | 0.00098 | 0.0004-0.0023 |
| *Pseudaphritis urvillii* | 0.00079 | 0.0005-0.0011 |
| Eleginopsioidea | 0.00228 | 0.0010-0.0039 |
| *Eleginops maclovinus* | 0.00168 | 0.0009-0.0024 |
| Cryonotothenioidea | 0.00218 | 0.0008-0.0038 |
| *Dissostichus mawsoni* | 0.00042 | 0.0001-0.0009 |
| *Trematomus scottii* | 0.00061 | 0.0002-0.0010 |
| *Gobionotothen gibberifrons* | 0.00078 | 0.0004-0.0012 |
| *Notothenia coriiceps* | 0.00076 | 0.0005-0.0012 |
| *Pogonophryne scotti* | 0.00071 | 0.0004-0.0011 |
| *Harpagifer antarcticus* | 0.00076 | 0.0005-0.0012 |
| *Chaenocephalus aceratus* | 0.00092 | 0.0003-0.0022 |
| *Bathydraco marri* | 0.00069 | 0.0003-0.0011 |

### Table S5. Positive selection in bone-associated† genes in Eleginopsioidea

| Ensembl Ortholog | Gene | P | P (Bonferroni) | LRT |
|---|---|---|---|---|
| ENSGACG00000014066 | LRRK1 | 1.46E-03 | 2.93E-03 | 10.922 |
| ENSGACG00000001740 | prdm1a | 5.90E-03 | 1.18E-02 | 8.170 |
| ENSGACG00000014774 | cahz | 4.37E-02 | 8.73E-02 | 4.251 |
| ENSGACG00000011819 | col12a1b | 4.44E-02 | 8.88E-02 | 4.218 |
| ENSGACG00000010282 | minpp1b | 1.11E-02 | 2.21E-02 | 6.935 |
| ENSGACG00000013609 | tacr1a | 3.90E-02 | 7.81E-02 | 4.469 |
| ENSGACG00000016847 | sparc | 1.69E-03 | 3.39E-03 | 10.635 |
| ENSGACG00000013381 | GABBR1 | 2.69E-02 | 5.38E-02 | 5.192 |
| ENSGACG00000014923 | col5a2b | 1.25E-05 | 2.49E-05 | 20.389 |
| ENSGACG00000008814 | tmem119a | 2.14E-02 | 4.28E-02 | 5.640 |
| ENSGACG00000007303 | ctnnbip1 | 8.15E-03 | 1.63E-02 | 7.534 |
| ENSGACG00000018120 | IGFBP4 | 7.24E-05 | 1.45E-04 | 16.889 |
| ENSGACG00000005830 | prpsap2 | 2.48E-02 | 4.95E-02 | 5.354 |
| ENSGACG00000012442 | notch2 | 3.74E-02 | 7.47E-02 | 4.553 |
| ENSGACG00000013102 | mapk3 | 1.42E-03 | 2.83E-03 | 10.990 |
| ENSGACG00000003838 | si:ch211-285f17.1 | 8.50E-03 | 1.70E-02 | 7.451 |
| ENSGACG00000005488 | phex | 1.17E-02 | 2.34E-02 | 6.819 |
| ENSGACG00000004564 | hoxd10a | 2.44E-03 | 4.88E-03 | 9.911 |
| ENSGACG00000007087 | LIMD1 | 4.55E-03 | 9.11E-03 | 8.680 |
| ENSGACG00000006673 | rbp4 | 4.58E-04 | 9.17E-04 | 13.225 |
| ENSGACG00000017924 | ghrb | 2.67E-02 | 5.34E-02 | 5.208 |
| ENSGACG00000019885 | slc8a1a | 1.36E-02 | 2.72E-02 | 6.530 |
| ENSGACG00000011433 | sun1 | 8.17E-03 | 1.63E-02 | 7.529 |
| ENSGACG00000007343 | col9a2 | 2.79E-02 | 5.58E-02 | 5.120 |
| ENSGACG00000006287 | ufl1 | 1.88E-02 | 3.76E-02 | 5.892 |
| ENSGACG00000008639 | pkdcc | 1.49E-02 | 2.99E-02 | 6.343 |
| ENSGACG00000010609 | ptk2bb | 2.00E-02 | 4.00E-02 | 5.772 |
| ENSGACG00000010218 | gata1a | 1.51E-03 | 3.03E-03 | 10.856 |
| ENSGACG00000015542 | LOCENSGACG00000015542 | 8.65E-03 | 1.73E-02 | 7.416 |
| ENSGACG00000005143 | col1a1a | 5.37E-04 | 1.07E-03 | 12.912 |
| ENSGACG00000018619 | il6st | 3.97E-02 | 7.94E-02 | 4.435 |
| ENSGACG00000014732 | penka | 3.16E-02 | 6.33E-02 | 4.876 |
| ENSGACG00000016426 | LOCENSGACG00000016426 | 4.59E-02 | 9.18E-02 | 4.153 |
| ENSGACG00000011376 | plxnb1b | 1.07E-03 | 2.15E-03 | 11.537 |
| ENSGACG00000006690 | TNFRSF11B | 2.58E-02 | 5.15E-02 | 5.277 |

| | | | | |
|---|---|---|---|---|
| ENSGACG00000003940 | ufd1l | 3.28E-02 | 6.56E-02 | 4.807 |
| ENSGACG00000009412 | LOCENSGACG00000009412 | 1.94E-04 | 3.87E-04 | 14.935 |
| ENSGACG00000016814 | slc26a2 | 4.17E-03 | 8.34E-03 | 8.853 |
| ENSGACG00000012223 | trip11 | 1.35E-02 | 2.71E-02 | 6.537 |
| ENSGACG00000008029 | PCSK5 | 4.34E-02 | 8.68E-02 | 4.262 |
| ENSGACG00000011806 | LOCENSGACG00000011806 | 2.28E-02 | 4.57E-02 | 5.512 |
| ENSGACG00000004355 | EIF2AK2 | 6.96E-04 | 1.39E-03 | 12.397 |
| ENSGACG00000006819 | col2a1b | 1.50E-02 | 3.00E-02 | 6.337 |
| ENSGACG00000004632 | KLF10 | 4.31E-06 | 8.63E-06 | 22.504 |
| ENSGACG00000019023 | actn3a | 2.03E-02 | 4.06E-02 | 5.740 |
| ENSGACG00000010790 | six2a | 4.38E-03 | 8.76E-03 | 8.757 |
| ENSGACG00000014956 | eng1b | 2.25E-02 | 4.51E-02 | 5.539 |
| ENSGACG00000018018 | NPR2 | 9.96E-03 | 1.99E-02 | 7.139 |
| ENSGACG00000000260 | itgav | 1.89E-02 | 3.79E-02 | 5.877 |
| ENSGACG00000002837 | sulf1 | 1.84E-02 | 3.69E-02 | 5.930 |
| ENSGACG00000009713 | matn3a | 1.35E-02 | 2.69E-02 | 6.548 |
| ENSGACG00000006853 | usp1 | 2.92E-02 | 5.83E-02 | 5.035 |
| ENSGACG00000006537 | mcph1 | 1.49E-02 | 2.97E-02 | 6.353 |
| ENSGACG00000017894 | spp1 | 2.59E-02 | 5.19E-02 | 5.264 |
| ENSGACG00000008948 | itgb3a | 1.01E-03 | 2.01E-03 | 11.666 |
| ENSGACG00000014564 | hdac4 | 3.97E-02 | 7.93E-02 | 4.437 |
| ENSGACG00000016287 | CSGALNACT1 | 3.48E-02 | 6.95E-02 | 4.694 |
| ENSGACG00000012231 | LOCENSGACG00000012231 | 4.13E-04 | 8.27E-04 | 13.430 |
| ENSGACG00000017458 | lef1 | 3.11E-02 | 6.22E-02 | 4.909 |
| ENSGACG00000018036 | slc34a1a | 4.85E-02 | 9.71E-02 | 4.046 |
| ENSGACG00000020663 | ryr1a | 1.72E-02 | 3.45E-02 | 6.064 |
| ENSGACG00000018814 | ano6 | 1.44E-02 | 2.88E-02 | 6.416 |
| ENSGACG00000016719 | col11a1b | 3.84E-03 | 7.69E-03 | 9.014 |
| ENSGACG00000004416 | eif2ak3 | 2.15E-02 | 4.30E-02 | 5.629 |
| ENSGACG00000015182 | NCAN (1 of many) | 9.69E-03 | 1.94E-02 | 7.193 |
| ENSGACG00000016060 | ddx21 | 2.79E-02 | 5.59E-02 | 5.119 |
| ENSGACG00000016773 | FBN1 | 1.15E-02 | 2.30E-02 | 6.857 |
| ENSGACG00000015511 | jund | 4.29E-02 | 8.59E-02 | 4.284 |
| ENSGACG00000012007 | mst1rb | 1.62E-02 | 3.24E-02 | 6.184 |
| ENSGACG00000004964 | nf1b | 3.70E-03 | 7.40E-03 | 9.090 |
| ENSGACG00000020221 | spns2 | 8.60E-03 | 1.72E-02 | 7.428 |
| ENSGACG00000011567 | LOCENSGACG00000011567 | 2.97E-02 | 5.94E-02 | 4.999 |
| ENSGACG00000003402 | foxp1a | 4.78E-02 | 9.57E-02 | 4.074 |
| ENSGACG00000008971 | isg15 | 2.29E-07 | 4.58E-07 | 28.358 |
| ENSGACG00000010238 | igsf10 | 3.86E-02 | 7.71E-02 | 4.491 |

| | | | | |
|---|---|---|---|---|
| ENSGACG00000016670 | mmp2 | 1.69E-02 | 3.38E-02 | 6.103 |
| ENSGACG00000008267 | nog2 | 1.56E-03 | 3.12E-03 | 10.795 |
| ENSGACG00000006511 | col1a2 | 7.06E-04 | 1.41E-03 | 12.367 |
| ENSORLG00000002037 | LOCENSORLG00000002037 | 5.35E-03 | 1.07E-02 | 8.363 |

†Select gene ontology IDs associated with bone development and mineralization (637 genes total)

GO:0001501,GO:0001503,GO:0001649,GO:0001957,GO:0001958,GO:0002051,GO:0002076
GO:0002158,GO:0003433,GO:0030278,GO:0030279,GO:0030282,GO:0030316,GO:0030500
GO:0030501,GO:0030502,GO:0031214,GO:0033687,GO:0033688,GO:0033689,GO:0033690
GO:0035630,GO:0036035,GO:0036179,GO:0043931,GO:0043932,GO:0044339,GO:0045667
GO:0045668,GO:0045669,GO:0045670,GO:0045671,GO:0045672,GO:0045778,GO:0048706
GO:0060351,GO:0070168,GO:0072674,GO:0072675,GO:0090290,GO:0090291,GO:1900158
GO:1900159

### Table S6. Accelerated sequence evolution in bone-associated† genes in Eleginopsioidea

| Ensembl Ortholog | Gene | P | P (Bonferroni) | LRT |
|---|---|---|---|---|
| ENSGACG00000014564 | hdac4 | 0.00E+00 | 0.00E+00 | 45.267 |
| ENSGACG00000002204 | fasn | 0.00E+00 | 0.00E+00 | 37.930 |
| ENSGACG00000003893 | shhb | 0.00E+00 | 0.00E+00 | 28.550 |
| ENSGACG00000003678 | zic1 | 0.00E+00 | 0.00E+00 | 21.956 |
| ENSGACG00000008942 | deaf1 | 0.00E+00 | 0.00E+00 | 20.953 |
| ENSGACG00000014956 | eng1b | 0.00E+00 | 0.00E+00 | 20.861 |
| ENSGACG00000003838 | si:ch211-285f17.1 | 0.00E+00 | 0.00E+00 | 18.399 |
| ENSGACG00000014897 | CHD7 | 0.00E+00 | 0.00E+00 | 17.048 |
| ENSGACG00000011376 | plxnb1b | 0.00E+00 | 0.00E+00 | 15.932 |
| ENSGACG00000017481 | gtpbp4 | 0.00E+00 | 0.00E+00 | 15.515 |
| ENSGACG00000003906 | npr3 | 0.00E+00 | 0.00E+00 | 14.273 |
| ENSGACG00000016083 | notch1a | 0.00E+00 | 0.00E+00 | 12.690 |
| ENSGACG00000020196 | wnt11r | 0.00E+00 | 0.00E+00 | 11.440 |
| ENSGACG00000020771 | zbtb16a | 1.00E-05 | 2.00E-05 | 9.754 |
| ENSGACG00000005297 | rarab | 1.00E-05 | 2.00E-05 | 9.003 |
| ENSGACG00000020663 | ryr1a | 1.00E-05 | 2.00E-05 | 8.860 |
| ENSGACG00000016835 | nipblb | 5.00E-05 | 1.00E-04 | 7.520 |
| ENSGACG00000011665 | syncrip | 5.00E-05 | 1.00E-04 | 7.517 |
| ENSGACG00000007999 | RARB | 7.00E-05 | 1.40E-04 | 7.286 |
| ENSORLG00000009503 | chd | 1.70E-04 | 3.40E-04 | 6.433 |
| ENSGACG00000016814 | slc26a2 | 1.70E-04 | 3.40E-04 | 6.426 |
| ENSGACG00000005810 | ACVR1 | 2.30E-04 | 4.60E-04 | 6.129 |
| ENSGACG00000020010 | tmem178 | 2.50E-04 | 5.00E-04 | 6.047 |
| ENSGACG00000010046 | id1 | 3.10E-04 | 6.20E-04 | 5.863 |
| ENSGACG00000008814 | tmem119a | 4.00E-04 | 8.00E-04 | 5.626 |
| ENSGACG00000016310 | sh3pxd2aa | 4.50E-04 | 9.00E-04 | 5.520 |
| ENSGACG00000006853 | usp1 | 4.60E-04 | 9.20E-04 | 5.488 |
| ENSGACG00000016468 | foxc1a | 4.70E-04 | 9.40E-04 | 5.465 |
| ENSGACG00000005279 | gdf6a | 4.80E-04 | 9.60E-04 | 5.445 |
| ENSGACG00000008289 | trim45 | 7.10E-04 | 1.42E-03 | 5.088 |
| ENSGACG00000010282 | minpp1b | 8.50E-04 | 1.70E-03 | 4.924 |
| ENSGACG00000006287 | ufl1 | 9.50E-04 | 1.90E-03 | 4.820 |
| ENSGACG00000020147 | osr1 | 1.11E-03 | 2.22E-03 | 4.678 |
| ENSGACG00000018575 | hmgcs1 | 1.43E-03 | 2.86E-03 | 4.448 |
| ENSGACG00000005508 | sfrp1a | 1.46E-03 | 2.92E-03 | 4.425 |

| | | | | |
|---|---|---|---|---|
| ENSGACG00000010238 | igsf10 | 2.06E-03 | 4.12E-03 | 4.116 |
| ENSGACG00000004632 | KLF10 | 2.07E-03 | 4.14E-03 | 4.113 |
| ENSGACG00000012411 | SMOC1 | 2.15E-03 | 4.30E-03 | 4.076 |
| ENSGACG00000020486 | pafah1b1b | 2.18E-03 | 4.36E-03 | 4.066 |
| ENSGACG00000014186 | fgf3 | 2.31E-03 | 4.62E-03 | 4.010 |
| ENSGACG00000003402 | foxp1a | 2.45E-03 | 4.90E-03 | 3.960 |
| ENSGACG00000009557 | alx1 | 2.62E-03 | 5.24E-03 | 3.899 |
| ENSGACG00000004931 | dlx1a | 2.82E-03 | 5.64E-03 | 3.831 |
| ENSGACG00000004834 | gli3 | 3.34E-03 | 6.68E-03 | 3.680 |
| ENSGACG00000011493 | si:dkey-42p8.3 | 3.39E-03 | 6.78E-03 | 3.666 |
| ENSGACG00000014119 | acp2 | 3.63E-03 | 7.26E-03 | 3.604 |
| ENSGACG00000005488 | phex | 3.90E-03 | 7.80E-03 | 3.540 |
| ENSGACG00000007912 | smurf1 | 3.94E-03 | 7.88E-03 | 3.531 |
| ENSGACG00000012407 | ercc2 | 4.26E-03 | 8.52E-03 | 3.461 |
| ENSGACG00000004764 | DNAJC13 | 4.62E-03 | 9.24E-03 | 3.387 |
| ENSGACG00000015083 | acvr2ab | 4.80E-03 | 9.60E-03 | 3.354 |
| ENSGACG00000008948 | itgb3a | 4.96E-03 | 9.92E-03 | 3.325 |
| ENSGACG00000009396 | hoxc9a | 7.28E-03 | 1.46E-02 | 2.984 |
| ENSGACG00000007341 | ireb2 | 7.42E-03 | 1.48E-02 | 2.968 |
| ENSGACG00000009872 | kat7b | 7.75E-03 | 1.55E-02 | 2.930 |
| ENSGACG00000013817 | BMPR2 | 8.04E-03 | 1.61E-02 | 2.897 |
| ENSGACG00000012967 | MATN3 | 8.53E-03 | 1.71E-02 | 2.845 |
| ENSGACG00000012822 | LOCENSGACG00000012822 | 8.89E-03 | 1.78E-02 | 2.809 |
| ENSGACG00000007795 | inppl1a | 9.54E-03 | 1.91E-02 | 2.747 |
| ENSGACG00000004680 | sema7a | 9.61E-03 | 1.92E-02 | 2.741 |

†Select gene ontology IDs associated with bone development and mineralization (637 genes total)

GO:0001501,GO:0001503,GO:0001649,GO:0001957,GO:0001958,GO:0002051,GO:0002076
GO:0002158,GO:0003433,GO:0030278,GO:0030279,GO:0030282,GO:0030316,GO:0030500
GO:0030501,GO:0030502,GO:0031214,GO:0033687,GO:0033688,GO:0033689,GO:0033690
GO:0035630,GO:0036035,GO:0036179,GO:0043931,GO:0043932,GO:0044339,GO:0045667
GO:0045668,GO:0045669,GO:0045670,GO:0045671,GO:0045672,GO:0045778,GO:0048706
GO:0060351,GO:0070168,GO:0072674,GO:0072675,GO:0090290,GO:0090291,GO:1900158
GO:1900159

### Table S7. Positive selection in bone-associated[†] genes in Channichthyidae

| Ensembl Ortholog | Gene | P | P (Bonferroni) | LRT |
|---|---|---|---|---|
| ENSGACG00000014066 | LRRK1 | 0.00E+00 | 0.00E+00 | 101.842 |
| ENSGACG00000010790 | six2a | 2.65E-10 | 5.29E-10 | 41.853 |
| ENSGACG00000010565 | tgfb3 | 7.14E-06 | 1.43E-05 | 21.500 |
| ENSGACG00000016783 | tbx3a | 8.11E-06 | 1.62E-05 | 21.248 |
| ENSGACG00000017894 | spp1 | 2.50E-04 | 5.00E-04 | 14.429 |
| ENSGACG00000011014 | aak1a | 4.19E-04 | 8.38E-04 | 13.403 |
| ENSGACG00000005143 | col1a1a | 7.30E-04 | 1.46E-03 | 12.301 |
| ENSGACG00000012822 | LOCENSGACG00000012822 | 1.37E-03 | 2.75E-03 | 11.048 |
| ENSGACG00000007087 | LIMD1 | 1.85E-03 | 3.71E-03 | 10.455 |
| ENSGACG00000010042 | BCAP29 | 3.20E-03 | 6.40E-03 | 9.376 |
| ENSGACG00000016860 | mfge8a | 4.62E-03 | 9.23E-03 | 8.654 |
| ENSGACG00000017255 | inpp4b | 4.76E-03 | 9.52E-03 | 8.593 |
| ENSGACG00000012007 | mst1rb | 5.49E-03 | 1.10E-02 | 8.310 |
| ENSGACG00000016468 | foxc1a | 6.33E-03 | 1.27E-02 | 8.030 |
| ENSGACG00000013533 | tfa | 8.34E-03 | 1.67E-02 | 7.487 |
| ENSGACG00000000393 | il19l | 1.19E-02 | 2.39E-02 | 6.783 |
| ENSGACG00000003946 | gpm6bb | 1.24E-02 | 2.47E-02 | 6.714 |
| ENSGACG00000016773 | FBN1 | 1.33E-02 | 2.65E-02 | 6.577 |
| ENSGACG00000020771 | zbtb16a | 1.82E-02 | 3.64E-02 | 5.958 |
| ENSGACG00000011694 | axin2 | 1.95E-02 | 3.89E-02 | 5.824 |
| ENSGACG00000008971 | isg15 | 2.40E-02 | 4.80E-02 | 5.416 |
| ENSGACG00000011781 | ocstamp | 3.01E-02 | 6.02E-02 | 4.975 |
| ENSGACG00000009662 | NA | 3.13E-02 | 6.26E-02 | 4.898 |
| ENSORLG00000002037 | LOCENSORLG00000002037 | 3.85E-02 | 7.70E-02 | 4.495 |
| ENSGACG00000004134 | chad | 4.95E-02 | 9.89E-02 | 4.009 |

†Select gene ontology IDs associated with bone development and mineralization (637 genes total)

GO:0001501,GO:0001503,GO:0001649,GO:0001957,GO:0001958,GO:0002051,GO:0002076
GO:0002158,GO:0003433,GO:0030278,GO:0030279,GO:0030282,GO:0030316,GO:0030500
GO:0030501,GO:0030502,GO:0031214,GO:0033687,GO:0033688,GO:0033689,GO:0033690
GO:0035630,GO:0036035,GO:0036179,GO:0043931,GO:0043932,GO:0044339,GO:0045667
GO:0045668,GO:0045669,GO:0045670,GO:0045671,GO:0045672,GO:0045778,GO:0048706
GO:0060351,GO:0070168,GO:0072674,GO:0072675,GO:0090290,GO:0090291,GO:1900158
GO:1900159

Table S8. Accelerated sequence evolution in bone-associated† genes in Channichthyidae

| Ensembl Ortholog | Gene | P | P (Bonferroni) | LRT |
|---|---|---|---|---|
| ENSGACG00000018025 | gdf2 | 0.00E+00 | 0.00E+00 | 44.466 |
| ENSGACG00000014066 | LRRK1 | 0.00E+00 | 0.00E+00 | 18.742 |
| ENSGACG00000010790 | six2a | 0.00E+00 | 0.00E+00 | 16.153 |
| ENSGACG00000020111 | BBX | 0.00E+00 | 0.00E+00 | 16.044 |
| ENSGACG00000012007 | mst1rb | 0.00E+00 | 0.00E+00 | 10.213 |
| ENSGACG00000012223 | trip11 | 1.00E-05 | 2.00E-05 | 9.635 |
| ENSGACG00000005143 | col1a1a | 1.00E-05 | 2.00E-05 | 9.225 |
| ENSGACG00000011839 | FOXP1 | 1.00E-05 | 2.00E-05 | 8.772 |
| ENSGACG00000003352 | sec22c | 2.00E-05 | 4.00E-05 | 8.542 |
| ENSGACG00000018575 | hmgcs1 | 4.00E-05 | 8.00E-05 | 7.720 |
| ENSGACG00000014838 | ptk2ba | 6.00E-05 | 1.20E-04 | 7.335 |
| ENSGACG00000018814 | ano6 | 7.00E-05 | 1.40E-04 | 7.308 |
| ENSGACG00000005830 | prpsap2 | 1.40E-04 | 2.80E-04 | 6.617 |
| ENSGACG00000016287 | CSGALNACT1 (2 of 2) | 1.60E-04 | 3.20E-04 | 6.447 |
| ENSGACG00000014536 | xpr1b | 1.70E-04 | 3.40E-04 | 6.423 |
| ENSGACG00000011843 | cdh11 | 6.30E-04 | 1.26E-03 | 5.194 |
| ENSGACG00000007087 | LIMD1 | 6.50E-04 | 1.30E-03 | 5.165 |
| ENSGACG00000014186 | fgf3 | 8.90E-04 | 1.78E-03 | 4.878 |
| ENSGACG00000018497 | AXIN2 (1 of many) | 1.16E-03 | 2.32E-03 | 4.640 |
| ENSGACG00000013007 | skia | 1.82E-03 | 3.64E-03 | 4.227 |
| ENSGACG00000016062 | col27a1a | 2.59E-03 | 5.18E-03 | 3.907 |
| ENSGACG00000016083 | notch1a | 2.80E-03 | 5.60E-03 | 3.838 |
| ENSGACG00000011349 | src | 3.32E-03 | 6.64E-03 | 3.684 |
| ENSGACG00000019404 | psmc2 | 3.69E-03 | 7.38E-03 | 3.589 |
| ENSGACG00000001547 | gli2a | 3.96E-03 | 7.92E-03 | 3.525 |
| ENSGACG00000014550 | txlng | 4.58E-03 | 9.16E-03 | 3.395 |
| ENSGACG00000007522 | jag2b | 6.46E-03 | 1.29E-02 | 3.090 |
| ENSGACG00000016016 | fam73b | 6.48E-03 | 1.30E-02 | 3.087 |

†Select gene ontology IDs associated with bone development and mineralization (637 genes total)

GO:0001501,GO:0001503,GO:0001649,GO:0001957,GO:0001958,GO:0002051,GO:0002076
GO:0002158,GO:0003433,GO:0030278,GO:0030279,GO:0030282,GO:0030316,GO:0030500
GO:0030501,GO:0030502,GO:0031214,GO:0033687,GO:0033688,GO:0033689,GO:0033690
GO:0035630,GO:0036035,GO:0036179,GO:0043931,GO:0043932,GO:0044339,GO:0045667
GO:0045668,GO:0045669,GO:0045670,GO:0045671,GO:0045672,GO:0045778,GO:0048706
GO:0060351,GO:0070168,GO:0072674,GO:0072675,GO:0090290,GO:0090291,GO:1900158
GO:1900159

#### Table S9. Tissue samples used in this study

| Family | Species | Source | ID# |
|---|---|---|---|
| Artedidraconidae | *Artedidraco skottsbergi* | No voucher | YFTC 4131 |
| Artedidraconidae | *Artedidraco skottsbergi* | No voucher | YFTC 4132 |
| Artedidraconidae | *Artedidraco skottsbergi* | YPM ICH 016389 | YFTC 7804 |
| Artedidraconidae | *Artedidraco skottsbergi* | YPM ICH 016389 | YFTC 7805 |
| Artedidraconidae | *Artedidraco skottsbergi* | YPM ICH 022479 | YFTC 15048 |
| Artedidraconidae | *Dolloidraco longedorsalis* | Detrich lab | HWD 53 |
| Artedidraconidae | *Dolloidraco longedorsalis* | Detrich lab | HWD 54 |
| Artedidraconidae | *Dolloidraco longedorsalis* | Detrich lab | HWD 55 |
| Artedidraconidae | *Dolloidraco longedorsalis* | Detrich lab | HWD 56 |
| Artedidraconidae | *Dolloidraco longedorsalis* | Detrich lab | HWD 57 |
| Artedidraconidae | *Histiodraco velifer* | No voucher | YFTC 2049 |
| Artedidraconidae | *Histiodraco velifer* | No voucher | YFTC 4135 |
| Artedidraconidae | *Histiodraco velifer* | No voucher | YFTC 4136 |
| Artedidraconidae | *Histiodraco velifer* | No voucher | YFTC 13801 |
| Artedidraconidae | *Histiodraco velifer* | No voucher | YFTC 14514 |
| Artedidraconidae | *Pogonophryne barsukovi* | YPM ICH 022357 | YFTC 15530 |
| Artedidraconidae | *Pogonophryne barsukovi* | YPM ICH 022357 | YFTC 15531 |
| Artedidraconidae | *Pogonophryne barsukovi* | YPM ICH 022357 | YFTC 15532 |
| Artedidraconidae | *Pogonophryne barsukovi* | YPM ICH 022357 | YFTC 15533 |
| Artedidraconidae | *Pogonophryne barsukovi* | YPM ICH 022357 | YFTC 15534 |
| Artedidraconidae | *Pogonophryne scotti* | YPM ICH 022357 | YFTC 15105 |
| Artedidraconidae | *Pogonophryne scotti* | YPM ICH 022553 | YFTC 15106 |
| Artedidraconidae | *Pogonophryne scotti* | YPM ICH 022553 | YFTC 15107 |
| Artedidraconidae | *Pogonophryne scotti* | YPM ICH 022553 | YFTC 15108 |
| Artedidraconidae | *Pogonophryne scotti* | YPM ICH 022553 | YFTC 15109 |
| Bathydraconidae | *Akarotaxis nudiceps* | YPM ICH 024046 | YFTC 20912 |
| Bathydraconidae | *Akarotaxis nudiceps* | YPM ICH 024238 | YFTC 20924 |
| Bathydraconidae | *Akarotaxis nudiceps* | YPM ICH 024118 | YFTC 20928 |
| Bathydraconidae | *Akarotaxis nudiceps* | YPM ICH 024118 | YFTC 20929 |
| Bathydraconidae | *Akarotaxis nudiceps* | YPM ICH 024118 | YFTC 20930 |
| Bathydraconidae | *Bathydraco marri* | NMNZ 043393 | YFTC 13884 |
| Bathydraconidae | *Bathydraco marri* | NMNZ 043394 | YFTC 13885 |
| Bathydraconidae | *Bathydraco marri* | NMNZ 043553 | YFTC 13886 |
| Bathydraconidae | *Bathydraco marri* | NMNZ 043634 | YFTC 13887 |
| Bathydraconidae | *Bathydraco marri* | NMNZ 043635 | YFTC 13888 |
| Bathydraconidae | *Gerlachea australis* | NMNZ 043338 | YFTC 13898 |
| Bathydraconidae | *Gerlachea australis* | NMNZ 043495 | YFTC 13899 |

| | | | |
|---|---|---|---|
| Bathydraconidae | *Gerlachea australis* | NMNZ 043517 | YFTC 13900 |
| Bathydraconidae | *Gerlachea australis* | NMNZ 043518 | YFTC 13901 |
| Bathydraconidae | *Gerlachea australis* | NMNZ 043545 | YFTC 13902 |
| Bathydraconidae | *Gymnodraco acuticeps* | Detrich lab | HWD 6 |
| Bathydraconidae | *Gymnodraco acuticeps* | Detrich lab | HWD 7 |
| Bathydraconidae | *Gymnodraco acuticeps* | Detrich lab | HWD 8 |
| Bathydraconidae | *Gymnodraco acuticeps* | Detrich lab | HWD 9 |
| Bathydraconidae | *Gymnodraco acuticeps* | Detrich lab | HWD 10 |
| Bathydraconidae | *Gymnodraco acuticeps* | Detrich lab | HWD 11 |
| Bathydraconidae | *Parachaenichthys charcoti* | Detrich lab | HWD 24 |
| Bathydraconidae | *Parachaenichthys charcoti* | Detrich lab | HWD 25 |
| Bathydraconidae | *Parachaenichthys charcoti* | Detrich lab | HWD 26 |
| Bathydraconidae | *Vomeridens infuscipinnis* | YPM ICH 020008 | YFTC 12880 |
| Bathydraconidae | *Vomeridens infuscipinnis* | YPM ICH 020009 | YFTC 12881 |
| Bathydraconidae | *Vomeridens infuscipinnis* | YPM ICH 020041 | YFTC 12884 |
| Bathydraconidae | *Vomeridens infuscipinnis* | YPM ICH 020025 | YFTC 12882 |
| Bathydraconidae | *Vomeridens infuscipinnis* | YPM ICH 020061 | YFTC 12919 |
| Bovichtidae | *Bovichtus diacanthus* | YPM ICH 021534 | YFTC 3482 |
| Bovichtidae | *Bovichtus diacanthus* | YPM ICH 021534 | YFTC 3483 |
| Bovichtidae | *Bovichtus diacanthus* | YPM ICH 021534 | YFTC 3484 |
| Bovichtidae | *Bovichtus diacanthus* | YPM ICH 021534 | YFTC 3485 |
| Bovichtidae | *Bovichtus diacanthus* | YPM ICH 021534 | YFTC 3486 |
| Bovichtidae | *Cottoperca trigloides* | Detrich lab | HWD 71 |
| Bovichtidae | *Cottoperca trigloides* | Detrich lab | HWD 72 |
| Bovichtidae | *Cottoperca trigloides* | Detrich lab | HWD 73 |
| Bovichtidae | *Cottoperca trigloides* | Detrich lab | HWD 74 |
| Bovichtidae | *Cottoperca trigloides* | Detrich lab | HWD 75 |
| Channichthyidae | *Chaenocephalus aceratus* | Detrich lab | HWD 1 |
| Channichthyidae | *Chaenocephalus aceratus* | Detrich lab | HWD 2 |
| Channichthyidae | *Chaenocephalus aceratus* | Detrich lab | HWD 3 |
| Channichthyidae | *Chaenocephalus aceratus* | Detrich lab | HWD 4 |
| Channichthyidae | *Chaenocephalus aceratus* | Detrich lab | HWD 5 |
| Channichthyidae | *Chaenodraco wilsoni* | Detrich lab | HWD 76 |
| Channichthyidae | *Chaenodraco wilsoni* | Detrich lab | HWD 77 |
| Channichthyidae | *Chaenodraco wilsoni* | Detrich lab | HWD 78 |
| Channichthyidae | *Chaenodraco wilsoni* | Detrich lab | HWD 79 |
| Channichthyidae | *Chaenodraco wilsoni* | Detrich lab | HWD 80 |
| Channichthyidae | *Champsocephalus esox* | Detrich lab | HWD 41 |
| Channichthyidae | *Champsocephalus esox* | Detrich lab | HWD 42 |
| Channichthyidae | *Champsocephalus gunnari* | YPM ICH 022626 | YFTC 14750 |

| | | | |
|---|---|---|---|
| Channichthyidae | *Champsocephalus gunnari* | YPM ICH 022626 | YFTC 14751 |
| Channichthyidae | *Champsocephalus gunnari* | YPM ICH 022626 | YFTC 14752 |
| Channichthyidae | *Champsocephalus gunnari* | YPM ICH 022626 | YFTC 14753 |
| Channichthyidae | *Champsocephalus gunnari* | YPM ICH 022626 | YFTC 14754 |
| Channichthyidae | *Chionobathyscus dewitti* | Detrich lab | HWD 32 |
| Channichthyidae | *Chionobathyscus dewitti* | Detrich lab | HWD 33 |
| Channichthyidae | *Chionobathyscus dewitti* | Detrich lab | HWD 34 |
| Channichthyidae | *Chionobathyscus dewitti* | Detrich lab | HWD 35 |
| Channichthyidae | *Cryodraco antarcticus* | No voucher | YFTC 11045 |
| Channichthyidae | *Cryodraco antarcticus* | No voucher | YFTC 11046 |
| Channichthyidae | *Cryodraco antarcticus* | No voucher | YFTC 11047 |
| Channichthyidae | *Cryodraco antarcticus* | Detrich lab | HWD 81 |
| Channichthyidae | *Cryodraco antarcticus* | Detrich lab | HWD 82 |
| Channichthyidae | *Cryodraco antarcticus* | Detrich lab | HWD 83 |
| Channichthyidae | *Dacodraco hunteri* | NMNZ 043399 | YFTC 16112 |
| Channichthyidae | *Dacodraco hunteri* | NMNZ 043430 | YFTC 16113 |
| Channichthyidae | *Dacodraco hunteri* | NMNZ 043450 | YFTC 16114 |
| Channichthyidae | *Dacodraco hunteri* | NMNZ 043496 | YFTC 16115 |
| Channichthyidae | *Neopagetopsis ionah* | NMNZ 043314 | YFTC 16116 |
| Channichthyidae | *Neopagetopsis ionah* | NMNZ 043315 | YFTC 16117 |
| Channichthyidae | *Neopagetopsis ionah* | NMNZ 043316 | YFTC 16118 |
| Channichthyidae | *Neopagetopsis ionah* | NMNZ 043358 | YFTC 16119 |
| Channichthyidae | *Neopagetopsis ionah* | NMNZ 043378 | YFTC 16120 |
| Channichthyidae | *Pagetopsis macropterus* | YPM ICH 016490 | YFTC 7775 |
| Channichthyidae | *Pagetopsis macropterus* | YPM ICH 016490 | YFTC 7776 |
| Channichthyidae | *Pagetopsis macropterus* | YPM ICH 016490 | YFTC 7777 |
| Channichthyidae | *Pagetopsis macropterus* | YPM ICH 016490 | YFTC 7778 |
| Channichthyidae | *Pagetopsis macropterus* | YPM ICH 016490 | YFTC 7779 |
| Channichthyidae | *Pseudochaenichthys georgianus* | Detrich lab | HWD 43 |
| Channichthyidae | *Pseudochaenichthys georgianus* | Detrich lab | HWD 44 |
| Channichthyidae | *Pseudochaenichthys georgianus* | Detrich lab | HWD 45 |
| Channichthyidae | *Pseudochaenichthys georgianus* | Detrich lab | HWD 46 |
| Channichthyidae | *Pseudochaenichthys georgianus* | Detrich lab | HWD 47 |
| Eleginopsidae | *Eleginops maclovinus* | Detrich lab | HWD 58 |
| Eleginopsidae | *Eleginops maclovinus* | Detrich lab | HWD 59 |
| Eleginopsidae | *Eleginops maclovinus* | Detrich lab | HWD 60 |
| Eleginopsidae | *Eleginops maclovinus* | Detrich lab | HWD 61 |
| Eleginopsidae | *Eleginops maclovinus* | Detrich lab | HWD 62 |
| Eleginopsidae | *Eleginops maclovinus* | Detrich lab | HWD 63 |
| Harpagiferidae | *Harpagifer antarcticus* | YPM ICH 016640 | YFTC 4399 |

| | | | |
|---|---|---|---|
| Harpagiferidae | *Harpagifer antarcticus* | YPM ICH 016640 | YFTC 4400 |
| Harpagiferidae | *Harpagifer antarcticus* | YPM ICH 016640 | YFTC 4401 |
| Harpagiferidae | *Harpagifer antarcticus* | YPM ICH 016640 | YFTC 4402 |
| Harpagiferidae | *Harpagifer antarcticus* | YPM ICH 016640 | YFTC 4403 |
| Nototheniidae | *Aethotaxis mitopteryx* | YPM ICH 022552 | YFTC 15291 |
| Nototheniidae | *Aethotaxis mitopteryx* | YPM ICH 022552 | YFTC 15292 |
| Nototheniidae | *Aethotaxis mitopteryx* | YPM ICH 022552 | YFTC 15293 |
| Nototheniidae | *Aethotaxis mitopteryx* | YPM ICH 022552 | YFTC 15294 |
| Nototheniidae | *Aethotaxis mitopteryx* | YPM ICH 022552 | YFTC 15295 |
| Nototheniidae | *Dissostichus eleginoides* | Detrich lab | HWD 36 |
| Nototheniidae | *Dissostichus eleginoides* | Detrich lab | HWD 37 |
| Nototheniidae | *Dissostichus eleginoides* | Detrich lab | HWD 38 |
| Nototheniidae | *Dissostichus eleginoides* | Detrich lab | HWD 39 |
| Nototheniidae | *Dissostichus eleginoides* | Detrich lab | HWD 40 |
| Nototheniidae | *Dissostichus mawsoni* | YPM ICH 022349 | YFTC 15462 |
| Nototheniidae | *Dissostichus mawsoni* | YPM ICH 022397 | YFTC 15346 |
| Nototheniidae | *Dissostichus mawsoni* | YPM ICH 022397 | YFTC 15347 |
| Nototheniidae | *Dissostichus mawsoni* | YPM ICH 022397 | YFTC 15348 |
| Nototheniidae | *Dissostichus mawsoni* | YPM ICH 022397 | YFTC 15349 |
| Nototheniidae | *Gobionotothen gibberifrons* | No voucher | YFTC 12759 |
| Nototheniidae | *Gobionotothen gibberifrons* | No voucher | YFTC 12760 |
| Nototheniidae | *Gobionotothen gibberifrons* | No voucher | YFTC 12761 |
| Nototheniidae | *Gobionotothen gibberifrons* | No voucher | YFTC 12762 |
| Nototheniidae | *Gobionotothen gibberifrons* | No voucher | YFTC 12763 |
| Nototheniidae | *Nototheniops nudifrons* | YPM ICH 020081 | YFTC 3809 |
| Nototheniidae | *Nototheniops nudifrons* | YPM ICH 020081 | YFTC 3810 |
| Nototheniidae | *Nototheniops nudifrons* | YPM ICH 020081 | YFTC 3811 |
| Nototheniidae | *Nototheniops nudifrons* | YPM ICH 020081 | YFTC 3812 |
| Nototheniidae | *Nototheniops nudifrons* | YPM ICH 020081 | YFTC 3813 |
| Nototheniidae | *Lepidonotothen squamifrons* | YPM ICH 028210 | YFTC 14785 |
| Nototheniidae | *Lepidonotothen squamifrons* | YPM ICH 028210 | YFTC 14786 |
| Nototheniidae | *Lepidonotothen squamifrons* | YPM ICH 022548 | YFTC 15065 |
| Nototheniidae | *Lepidonotothen squamifrons* | YPM ICH 022548 | YFTC 15066 |
| Nototheniidae | *Lepidonotothen squamifrons* | YPM ICH 022548 | YFTC 15067 |
| Nototheniidae | *Paranotothenia angustata* | Detrich lab | HWD 12 |
| Nototheniidae | *Paranotothenia angustata* | Detrich lab | HWD 13 |
| Nototheniidae | *Paranotothenia angustata* | Detrich lab | HWD 14 |
| Nototheniidae | *Paranotothenia angustata* | Detrich lab | HWD 15 |
| Nototheniidae | *Paranotothenia angustata* | Detrich lab | HWD 16 |
| Nototheniidae | *Notothenia coriiceps* | Detrich lab | HWD 27 |

| | | | |
|---|---|---|---|
| Nototheniidae | *Notothenia coriiceps* | Detrich lab | HWD 28 |
| Nototheniidae | *Notothenia coriiceps* | Detrich lab | HWD 29 |
| Nototheniidae | *Notothenia coriiceps* | Detrich lab | HWD 30 |
| Nototheniidae | *Notothenia coriiceps* | Detrich lab | HWD 31 |
| Nototheniidae | *Notothenia rossii* | Detrich lab | HWD 64 |
| Nototheniidae | *Notothenia rossii* | Detrich lab | HWD 65 |
| Nototheniidae | *Notothenia rossii* | Detrich lab | HWD 66 |
| Nototheniidae | *Notothenia rossii* | Detrich lab | HWD 67 |
| Nototheniidae | *Notothenia rossii* | Detrich lab | HWD 68 |
| Nototheniidae | *Notothenia rossii* | Detrich lab | HWD 69 |
| Nototheniidae | *Notothenia rossii* | Detrich lab | HWD 70 |
| Nototheniidae | *Trematomus borchgrevinki* | No voucher | YFTC 12070 |
| Nototheniidae | *Trematomus borchgrevinki* | No voucher | YFTC 12071 |
| Nototheniidae | *Trematomus borchgrevinki* | No voucher | YFTC 12072 |
| Nototheniidae | *Trematomus borchgrevinki* | No voucher | YFTC 13638 |
| Nototheniidae | *Patagonotothen squamiceps* | No voucher | YFTC 24046 |
| Nototheniidae | *Patagonotothen squamiceps* | No voucher | YFTC 24047 |
| Nototheniidae | *Patagonotothen squamiceps* | No voucher | YFTC 24048 |
| Nototheniidae | *Patagonotothen squamiceps* | No voucher | YFTC 24049 |
| Nototheniidae | *Patagonotothen squamiceps* | No voucher | YFTC 24050 |
| Nototheniidae | *Patagonotothen guntheri* | No voucher | YFTC 24057 |
| Nototheniidae | *Patagonotothen guntheri* | No voucher | YFTC 24058 |
| Nototheniidae | *Patagonotothen guntheri* | No voucher | YFTC 24059 |
| Nototheniidae | *Patagonotothen guntheri* | No voucher | YFTC 24060 |
| Nototheniidae | *Patagonotothen guntheri* | No voucher | YFTC 24061 |
| Nototheniidae | *Pleuragramma antarctica* | YPM ICH 022586 | YFTC 15002 |
| Nototheniidae | *Pleuragramma antarctica* | YPM ICH 022586 | YFTC 15003 |
| Nototheniidae | *Pleuragramma antarctica* | YPM ICH 022586 | YFTC 15004 |
| Nototheniidae | *Pleuragramma antarctica* | YPM ICH 022586 | YFTC 15005 |
| Nototheniidae | *Pleuragramma antarctica* | YPM ICH 022586 | YFTC 15006 |
| Nototheniidae | *Trematomus bernacchii* | Detrich lab | HWD 17 |
| Nototheniidae | *Trematomus bernacchii* | Detrich lab | HWD 18 |
| Nototheniidae | *Trematomus bernacchii* | Detrich lab | HWD 19 |
| Nototheniidae | *Trematomus bernacchii* | Detrich lab | HWD 20 |
| Nototheniidae | *Trematomus bernacchii* | Detrich lab | HWD 21 |
| Nototheniidae | *Trematomus bernacchii* | Detrich lab | HWD 22 |
| Nototheniidae | *Trematomus bernacchii* | Detrich lab | HWD 23 |
| Nototheniidae | *Trematomus eulepidotus* | YPM ICH 022564 | YFTC 15403 |
| Nototheniidae | *Trematomus eulepidotus* | YPM ICH 022564 | YFTC 15404 |
| Nototheniidae | *Trematomus eulepidotus* | YPM ICH 022564 | YFTC 15405 |

| | | | |
|---|---|---|---|
| Nototheniidae | *Trematomus eulepidotus* | YPM ICH 022564 | YFTC 15406 |
| Nototheniidae | *Trematomus eulepidotus* | YPM ICH 022564 | YFTC 15407 |
| Nototheniidae | *Trematomus hansoni* | Detrich lab | HWD 48 |
| Nototheniidae | *Trematomus hansoni* | Detrich lab | HWD 49 |
| Nototheniidae | *Trematomus hansoni* | Detrich lab | HWD 50 |
| Nototheniidae | *Trematomus hansoni* | Detrich lab | HWD 51 |
| Nototheniidae | *Trematomus hansoni* | Detrich lab | HWD 52 |
| Nototheniidae | *Trematomus newnesi* | YPM ICH 016464 | YFTC 7708 |
| Nototheniidae | *Trematomus newnesi* | YPM ICH 016464 | YFTC 7709 |
| Nototheniidae | *Trematomus newnesi* | YPM ICH 016464 | YFTC 7710 |
| Nototheniidae | *Trematomus newnesi* | YPM ICH 016464 | YFTC 7711 |
| Nototheniidae | *Trematomus newnesi* | YPM ICH 016464 | YFTC 7712 |
| Nototheniidae | *Trematomus scotti* | YPM ICH 024115 | YFTC 20940 |
| Nototheniidae | *Trematomus scotti* | YPM ICH 024115 | YFTC 20941 |
| Nototheniidae | *Trematomus scotti* | YPM ICH 024115 | YFTC 20942 |
| Nototheniidae | *Trematomus scotti* | YPM ICH 024115 | YFTC 20943 |
| Nototheniidae | *Trematomus scotti* | YPM ICH 024115 | YFTC 20944 |
| Percidae | *Percina caprodes* | YPM ICH 027375 | YFTC 24610 |
| Percidae | *Percina caprodes* | YPM ICH 027375 | YFTC 24610 |
| Percidae | *Percina caprodes* | YPM ICH 027375 | YFTC 24611 |
| Percidae | *Percina caprodes* | YPM ICH 027375 | YFTC 24612 |
| Percidae | *Percina caprodes* | YPM ICH 027375 | YFTC 24613 |
| Percophidae | *Percophis brasiliensis* | LBP 8668 | 35303 |
| Percophidae | *Percophis brasiliensis* | LBP 8654 | 35304 |
| Percophidae | *Percophis brasiliensis* | LBP 8654 | 35305 |
| Percophidae | *Percophis brasiliensis* | LBP 8668 | 35307 |
| Percophidae | *Percophis brasiliensis* | LBP 19527 | 73365 |
| Percophidae | *Percophis brasiliensis* | LBP 19527 | 73366 |
| Percophidae | *Percophis brasiliensis* | LBP 19527 | 73367 |
| Pseudaphritidae | *Pseudaphritis urvillii* | CSIRO 6892-01 | YFTC 16619 |
| Pseudaphritidae | *Pseudaphritis urvillii* | CSIRO 6892-01 | YFTC 16620 |
| Pseudaphritidae | *Pseudaphritis urvillii* | CSIRO 7637-05 | YFTC 16621 |

#### Table S10. Enrichment of genes with missing coverage in dataset

| GOTerm | Term Name | GO type | # genes | # genes low cov. † | fold enrichment | p-value | FDR qvalue |
|---|---|---|---|---|---|---|---|
| GO:0043410 | positive regulation of MAPK cascade | bp | 92 | 15 | 4.02 | 4.22E-06 | 1.72E-03 |
| GO:0030198 | extracellular matrix organization | bp | 209 | 30 | 3.54 | 1.89E-09 | 5.41E-06 |
| GO:0016925 | protein sumoylation | bp | 111 | 15 | 3.33 | 4.27E-05 | 1.36E-02 |
| GO:0006955 | immune response | bp | 268 | 29 | 2.67 | 1.70E-06 | 8.07E-04 |
| GO:0006954 | inflammatory response | bp | 308 | 33 | 2.64 | 4.19E-07 | 2.99E-04 |
| GO:0007166 | cell surface receptor signaling pathway | bp | 297 | 31 | 2.57 | 1.63E-06 | 8.07E-04 |
| GO:0018108 | peptidyl-tyrosine phosphorylation | bp | 211 | 21 | 2.45 | 1.47E-04 | 3.81E-02 |
| GO:0002376 | immune system process | bp | 251 | 24 | 2.36 | 9.61E-05 | 2.75E-02 |
| GO:0016032 | viral process | bp | 377 | 35 | 2.29 | 5.23E-06 | 1.87E-03 |
| GO:0007155 | cell adhesion | bp | 615 | 53 | 2.12 | 2.36E-07 | 2.25E-04 |
| GO:0016504 | peptidase activator activity | mf | 13 | 5 | 9.48 | 1.07E-04 | 1.01E-02 |
| GO:0005031 | tumor necrosis factor-activated receptor activity | mf | 23 | 6 | 6.43 | 2.45E-04 | 2.08E-02 |
| GO:0005201 | extracellular matrix structural constituent | mf | 86 | 21 | 6.02 | 2.27E-11 | 9.65E-09 |
| GO:0004714 | transmembrane receptor protein tyrosine kinase activity | mf | 89 | 13 | 3.60 | 6.03E-05 | 6.41E-03 |
| GO:0005198 | structural molecule activity | mf | 230 | 21 | 2.25 | 4.76E-04 | 3.38E-02 |
| GO:0003779 | actin binding | mf | 464 | 38 | 2.02 | 3.64E-05 | 4.43E-03 |
| GO:0000800 | lateral element | cc | 13 | 4 | 7.58 | 1.44E-03 | 2.48E-02 |
| GO:0005771 | multivesicular body | cc | 22 | 6 | 6.72 | 1.87E-04 | 5.02E-03 |
| GO:0005640 | nuclear outer membrane | cc | 22 | 5 | 5.60 | 1.61E-03 | 2.70E-02 |
| GO:0005581 | collagen trimer | cc | 111 | 18 | 4.00 | 5.16E-07 | 2.51E-05 |
| GO:0005604 | basement membrane | cc | 92 | 14 | 3.75 | 1.98E-05 | 6.23E-04 |
| GO:0030175 | filopodium | cc | 69 | 10 | 3.57 | 4.47E-04 | 9.58E-03 |
| GO:0005578 | proteinaceous extracellular matrix | cc | 371 | 46 | 3.06 | 1.80E-11 | 2.41E-09 |
| GO:0005814 | centriole | cc | 107 | 13 | 2.99 | 3.97E-04 | 8.87E-03 |
| GO:0072562 | blood microparticle | cc | 91 | 11 | 2.98 | 1.13E-03 | 2.09E-02 |
| GO:0009897 | external side of plasma membrane | cc | 166 | 20 | 2.97 | 1.40E-05 | 5.01E-04 |
| GO:0005923 | bicellular tight junction | cc | 158 | 18 | 2.81 | 7.78E-05 | 2.32E-03 |
| GO:0031225 | anchored component of membrane | cc | 88 | 10 | 2.80 | 2.98E-03 | 4.20E-02 |

| | | | | | | | |
|---|---|---|---|---|---|---|---|
| GO:0043235 | receptor complex | cc | 116 | 13 | 2.76 | 8.64E-04 | 1.78E-02 |
| GO:0005911 | cell-cell junction | cc | 193 | 19 | 2.43 | 3.43E-04 | 7.98E-03 |
| GO:0031012 | extracellular matrix | cc | 322 | 31 | 2.37 | 8.63E-06 | 3.30E-04 |
| GO:0005788 | endoplasmic reticulum lumen | cc | 194 | 17 | 2.16 | 2.45E-03 | 3.75E-02 |
| GO:0009986 | cell surface | cc | 498 | 42 | 2.08 | 7.11E-06 | 2.93E-04 |
| GO:0005874 | microtubule | cc | 357 | 29 | 2.00 | 3.35E-04 | 7.98E-03 |

bp – biological process, mf – molecular function, cc – cellular component

Table S11. Notothenioid specimens for CT scan

| Species | Catalog Number | Specimen # | STL (cm) |
|---|---|---|---|
| Bovichtus variegatus | YPM ICH 005976 | 5976 | 13.5 |
| Eleginops maclovinus | YPM ICH 009315 | NA | 32.0 |
| Dissostichus mawsoni | YPM ICH 020040 | 12848 | 27.5 |
| Trematomus bernacchii | YPM ICH 020469 | 20469 | 19.5 |
| Gobionotothen gibberifrons | YPM ICH 024029 | 20745 | 19.0 |
| Gymnodraco acuticeps | YPM ICH 022337 | 15454 | 24.0 |
| Champsocephalus gunnari | YPM ICH 016478 | 07959 | 31.0 |

#### Supplemental References


1. S. C. Shin *et al.*, The genome sequence of the Antarctic bullhead notothen reveals evolutionary adaptations to a cold environment. *Genome Biol.* **15**, 468 (2014).
2. M. Tine *et al.*, European sea bass genome and its variation provide insights into adaptation to euryhalinity and speciation. *Nat. Commun.* **5**, 5770 (2014).
3. J. Herrero *et al.*, Ensembl comparative genomics resources. *Database*. **2016**, 1–17 (2016).
4. A. Kozomara, S. Griffiths-Jones, MiRBase: Integrating microRNA annotation and deep-sequencing data. *Nucleic Acids Res.* **39**, D152-7 (2011).
5. S. Dimitrieva, P. Bucher, UCNEbase - A database of ultraconserved non-coding elements and genomic regulatory blocks. *Nucleic Acids Res.* **41**, 101–109 (2013).
6. A. R. Quinlan, I. M. Hall, BEDTools: a flexible suite of utilities for comparing genomic features. *Bioinformatics*. **26**, 841–2 (2010).
7. J. M. Daane, N. Rohner, P. Konstantinidis, S. Djuranovic, M. P. Harris, Parallelism and Epistasis in Skeletal Evolution Identified through Use of Phylogenomic Mapping Strategies. *Mol. Biol. Evol.* **33**, 1–12 (2016).
8. A. M. Bolger, M. Lohse, B. Usadel, Trimmomatic: A flexible trimmer for Illumina sequence data. *Bioinformatics*. **30**, 2114–2120 (2014).
9. S. Altschul, W. Gish, W. Miller, Basic Local Alignment Search Tool. *J Mol Biol.* **215**, 403–410 (1990).
10. X. Huang, CAP3: A DNA Sequence Assembly Program. *Genome Res.* **9**, 868–877 (1999).
11. F. J. Sedlazeck, P. Rescheneder, A. Von Haeseler, NextGenMap: Fast and accurate read mapping in highly polymorphic genomes. *Bioinformatics*. **29**, 2790–2791 (2013).
12. K. Katoh, K. Kuma, H. Toh, T. Miyata, MAFFT version 5: improvement in accuracy of multiple sequence alignment. *Nucleic Acids Res.* **33**, 511–8 (2005).
13. H. Li *et al.*, The Sequence Alignment/Map format and SAMtools. *Bioinformatics*. **25**, 2078–9 (2009).
14. J. T. Robinson *et al.*, Integrative Genome Viewer. *Nat. Biotechnol.* **29**, 24–6 (2011).
15. L. T. Nguyen, H. A. Schmidt, A. Von Haeseler, B. Q. Minh, IQ-TREE: A fast and effective stochastic algorithm for estimating maximum-likelihood phylogenies. *Mol. Biol. Evol.* **32**, 268–274 (2015).
16. K. Chen, D. Durand, M. Farach-Colton, NOTUNG: A Program for Dating Gene Duplications and Optimizing Gene Family Trees. *J. Comput. Biol.* **7**, 429–447 (2000).
17. V. Ranwez, S. Harispe, F. Delsuc, E. J. P. Douzery, MACSE: Multiple alignment of coding SEquences accounting for frameshifts and stop codons. *PLoS One*. **6** (2011).
18. I. Sela, H. Ashkenazy, K. Katoh, T. Pupko, GUIDANCE2: Accurate detection of unreliable alignment regions accounting for the uncertainty of multiple parameters. *Nucleic Acids Res.* **43**, W7–W14 (2015).
19. S. Kalyaanamoorthy, B. Q. Minh, T. K. F. Wong, A. Von Haeseler, L. S. Jermiin, ModelFinder: Fast model selection for accurate phylogenetic estimates. *Nat. Methods*. **14**, 587–589 (2017).
20. D. T. Hoang, O. Chernomor, A. Von Haeseler, B. Q. Minh, L. S. Vinh, UFBoot2: Improving the ultrafast bootstrap approximation. *Mol. Biol. Evol.* **35**, 518–522 (2018).
21. L. S. Kubatko, J. H. Degnan, Inconsistency of phylogenetic estimates from concatenated data under coalescence. *Syst. Biol.* **56**, 17–24 (2007).
22. S. Roch, M. Steel, Likelihood-based tree reconstruction on a concatenation of aligned

sequence data sets can be statistically inconsistent. *Theor. Popul. Biol.* **100**, 56–62 (2015).
23. C. Zhang, M. Rabiee, E. Sayyari, S. Mirarab, ASTRAL-III: Polynomial time species tree reconstruction from partially resolved gene trees. *BMC Bioinformatics*. **19**, 15–30 (2018).
24. E. Sayyari, S. Mirarab, Fast Coalescent-Based Computation of Local Branch Support from Quartet Frequencies. *Mol. Biol. Evol.* **33**, 1654–1668 (2016).
25. A. Dornburg, S. Federman, A. D. Lamb, C. D. Jones, T. J. Near, Cradles and museums of Antarctic teleost biodiversity. *Nat. Ecol. Evol.* (2017).
26. D. L. Rabosky *et al.*, BAMMtools: An R package for the analysis of evolutionary dynamics on phylogenetic trees. *Methods Ecol. Evol.* **5**, 701–707 (2014).
27. R. Bouckaert *et al.*, BEAST 2: A Software Platform for Bayesian Evolutionary Analysis. *PLoS Comput. Biol.* **10**, 1–6 (2014).
28. R. R. Bouckaert, A. J. Drummond, bModelTest: Bayesian phylogenetic site model averaging and model comparison. *BMC Evol. Biol.* **17**, 1–11 (2017).
29. A. J. Drummond, M. A. Suchard, Bayesian random local clocks, or one rate to rule them all. *BMC Biol.* **8**, 114 (2010).
30. T. J. Near *et al.*, Identification of the notothenioid sister lineage illuminates the biogeographic history of an Antarctic adaptive radiation. *BMC Evol. Biol.* **15**, 109 (2015).
31. M. D. Smith *et al.*, Less is more: An adaptive branch-site random effects model for efficient detection of episodic diversifying selection. *Mol. Biol. Evol.* **32**, 1342–1353 (2015).
32. S. L. Kosakovsky Pond, S. D. W. Frost, S. V. Muse, HyPhy: Hypothesis testing using phylogenies. *Bioinformatics*. **21**, 676–679 (2005).
33. K. S. Pollard, M. J. Hubisz, K. R. Rosenbloom, A. Siepel, Detection of nonneutral substitution rates on mammalian phylogenies. *Genome Res.* **20**, 110–21 (2010).
34. M. J. Hubisz, K. S. Pollard, A. Siepel, PHAST and RPHAST: phylogenetic analysis with space/time models. *Brief. Bioinform.* **12**, 41–51 (2011).
35. R. J. Kinsella *et al.*, Ensembl BioMarts: A hub for data retrieval across taxonomic space. *Database*. **2011**, 1–9 (2011).
36. S. Köhler *et al.*, The human phenotype ontology in 2017. *Nucleic Acids Res.* **45**, D865–D876 (2017).
37. J. T. Daub, S. Moretti, I. I. Davydov, L. Excoffier, Detection of Pathways Affected by Positive Selection in Primate Lineages Ancestral to Humans. *Mol. Biol. Evol.* **34**, 1391–1402 (2017).
38. C. Nüsslein-Volhard, R. Dahm, *Zebrafish: a practical approach* (Oxford University Press, 2002).
39. K. Henke *et al.*, Genetic Screen for Postembryonic Development in the Zebrafish (*Danio rerio*): Dominant Mutations Affecting Adult Form**,** *Genetics*, 207, 609-623 (2017).
40. T. J. Near, S. K. Parker, H. W. Detrich, A genomic fossil reveals key steps in hemoglobin loss by the Antarctic icefishes. *Mol. Biol. Evol.* **23**, 2008–2016 (2006).
